# Radiation therapy promotes unsaturated fatty acids to maintain survival of glioblastoma

**DOI:** 10.1101/2022.06.01.494338

**Authors:** Mara De Martino, Camille Daviaud, Hanna E. Minns, Nabeel Attarwala, Qiuying Chen, Noah Dephoure, Seung-Won Choi, Raùl Rabadàn, Robyn D. Gartrell, Evagelia C. Laiakis, Claire Vanpouille-Box

## Abstract

**Purpose:** Radiation therapy (RT) is essential for the management of glioblastoma (GBM). However, GBM frequently relapses within the irradiated margins, thus suggesting that RT might stimulate mechanisms of resistance that limits its efficacy. GBM is recognized for its metabolic plasticity, but whether RT-induced resistance relies on metabolic adaptation remains unclear.

**Methods:** We analyzed *in vitro* extracellular flux and profiled targeted metabolites as well as free fatty acids in two syngenic models of glioblastomas 24hrs post RT. Metabolic adaptation of irradiated GBM were confirmed *in vivo* by mass spectrometry imaging. The role of the fatty acid synthase (FASN) in RT-induced lipid metabolites was assessed by genetical and pharmacological inhibition of *Fasn* in irradiated GBM cells. The impact of FASN-mediated lipids on endoplasmic reticulum (ER) stress and apoptosis of irradiated GBM cells were performed by transmission electronic microscopy, western blot, clonogenic assay and flow cytometry. Inhibition of FASN combined with focal RT was assessed in mice. Analysis of a public dataset of GBM patients was performed to correlate preclinical findings.

**Results:** Here, we show *in vitro* and *in vivo* that irradiated GBM tumors switch their metabolic program to accumulate lipids, especially unsaturated fatty acids. This resulted in an increase formation of lipid droplets to prevent ER stress. We uncovered that FASN is critical for lipid accumulation of irradiated GBM and demonstrate that genetic suppression and pharmacological inhibition of FASN lead to mitochondrial dysfunction and apoptosis. Combination of FASN inhibition with focal RT improved the median survival of GBM-bearing mice. Supporting the translational value of these findings, retrospective analysis of the GLASS consortium dataset of matched GBM patients revealed an enrichment in lipid metabolism signature in recurrent GBM compared to primary.

**Conclusions:** Overall, these results demonstrate that RT drives GBM resistance by generating a lipogenic environment permissive to GBM survival. Targeting lipid metabolism might be required to develop more effective anti-GBM strategies.

## INTRODUCTION

Glioblastoma (GBM) is the most aggressive primary brain cancer in adults and accounts for 3-15% of central nervous system (CNS) tumors in children. A rising incidence of GBM has been observed over the last 15 years (1,2), while the median survival of GBM patients remains unchanged (15 months); consequently, GBM mortality is predicted to increase. Radiation therapy (RT) is the standard-of-care for the management of GBM (3,4) and is the most widely used treatment for inoperable brain tumors (5,6). Unfortunately, GBM recurrence often arises within the irradiated margins (7), suggesting that RT might not optimally control GBM and might generate and/or exacerbate resistance mechanisms to support GBM progression. The mechanisms by which RT drives evasion of GBM are unknown.

GBM is recognized for its altered tumor metabolism and for its ability to respond to metabolic stressors during cancer progression and treatment (8). For instance, GBM cells can metabolize glucose into lactate irrespective of the presence of oxygen to use glucose-derived carbons for the synthesis of essential cellular components, while generating enough ATP to support cellular processes (9). Consequently, while GBM (like many other tumors) relies on glycolysis as energy supply, it possesses the flexibility to switch metabolic programs in order to adapt to the heterogeneity of its tumor microenvironment (TME) which includes distinct regions of hypoxia, necrosis, angiogenesis and different availabilities of nutrients (10). In that context, other metabolic pathways such as glutamine and lipid metabolism are actively investigated as alternative energy sources to identify additional metabolic liabilities for anti-GBM therapies (11,12).

Oxidative stressors, like reactive oxygen species (ROS) generated by RT, have been shown to reprogram the metabolism of an irradiated tumor (11,13). Such metabolic reprogramming, in part, can promote the aggressivity of GBM. One study reported that a 20 Gy irradiation of healthy mouse brains prior to orthotopic implantation of human GBM cells worsens GBM outcome due to enhanced tumor cell proliferation and migration, as well as microglial activation (14). While increased levels of ATP and GTP with reduced levels of antioxidants, glutathione and ascorbate, were associated with the poor outcome of GBM in pre-irradiated brains (14), a comprehensive identification of the metabolic pathways altered in irradiated GBM tumors as well as their consequences for GBM growth and resistance is underexplored.

Here, we define the metabolic switch of GBM in response to RT *in vitro* and *in vivo*, uncovering the role of unsaturated fatty acids (UFAs) in providing metabolic plasticity of GBM; thereby allowing GBM cells to adapt to genotoxic stress and develop resistance mechanisms. This includes serving as a metabolic cue to prevent endoplasmic reticulum (ER) stress and resistance to apoptosis, while serving as an additional energy supply to permit survival of GBM. Specifically, our data demonstrate that accumulation of UFAs post RT leads to the formation of lipid droplets (LDs), a major site for the synthesis of the bioactive lipid prostaglandin E2 (PGE2) that is ascribed roles in cancer stemness and immunossupression. Analysis of the GLASS consortium dataset of GBM patients revealed an increased expression of lipid metabolism signatures as well as an increased enrichment score of the REACTOME fatty acid pathway in recurrent GBM compared to matched primary. Thus, supporting the clinical relevance of this regulatory mechanism.

Consequently, these observations indicate that RT induces a growth permissive metabolic reprogramming by increasing tumor-derived UFAs in GBM, suggesting that targeting lipid metabolism in the context of RT may be required to control GBM recurrence.

## METHODS AND MATERIALS

### Mice

Eight-week-old C57BL/6N female mice from Taconic Animal Laboratory (Germantown, NY) were maintained under pathogen-free conditions in the animal facility at Weill Cornell Medicine. All experiments were approved by the Institutional Animal Care and Use Committee.

### Cells and reagents

C57BL/6N mouse-derived GBM GL261 were obtained from S. Demaria (15) and CT2A were purchased from Millipore-Sigma (Burlington, MA). GL261 and CT2A are poorly immunogenic and share the growth characteristics as well as invasive properties of human GBM. Intracerebral (i.c.) injection of 5×10^4^ GL261 or CT2A cells into syngeneic C57BL/6N results in the death of the mice in approximately 20-25 days (15,16). U118 human GBM cells were purchased from ATCC (Manassas, VA). All cells were authenticated by IDEEX Bioresearch (Columbia, MO), and further characterized by morphology, phenotype, growth and pattern of metastasis *in vivo* and routinely screened for *Mycoplasma* (LookOut^®^ Mycoplasma PCR Detection kit, Sigma-Aldrich). None of the cells used are listed in the International Cell Line Authentication Committee (ICLAC) database of commonly misidentified cell lines. GL261, CT2A and U118 were cultured in DMEM (GIBCO) supplemented with 2 mmol/L L-glutamine, 100 U/mL penicillin, 100 μg/mL streptomycin, 2.5 × 10^−5^ mol/L 2-mercapthoethanol, 5 mM HEPES, 50 μg/mL gentamicin sulfate and 10% fetal bovine serum (FBS) (Life technologies). For experiments with doxycycline, cells were grown in media containing tetracycline-FBS and induced with 4μg/mL of doxycycline 10 days prior to treatment.

### shRNA-mediated gene knock-down and lentiviral transduction

HEK 293-FT cells were used to produce viruses upon transfection of the packaging plasmids pPAX2 and pMD2 and a pTRIPZ vector containing a tetracycline-inducible promoter (GE Dharmacon technology, kindly provided by Dr. Robert Schneider). ShRNAs directed against FASN mRNA (*Fasn*, mouse-shRNA: GACAGCGCATGCCTTTGTAAAC) or a non-silencing sequence (NS; mouse-human-shRNA: AATTCTCCGAACGTGTCACGT) were cloned into pTRIPZ using EcoRI and XhoI restriction sites. GL261 and CT2A cells were transduced with cellfree virus-containing supernatants and selected with 4μg/mL of puromycin for 48 hours.

### CRISPR/Cas9

To generate Fasn-deficient clones using CRISPR, GL261 and CT2A cells were transduced with lentiviral particles, to incorporate a lentiviral vector containing guide RNA (gRNA) and Cas9 coexpression (VectorBuilder, IL). Two *Fasn* targeting ccRNA were used (#1: pLV[CRISPR]- hCas9:T2A:Puro-U6>mFasn[gRNA#1485], ccRNA: GATGAACGTGTACCGGGACG; #2: pLV[CRISPR]-hCas9:T2A:Puro-U6>mFasn[gRNA#2246], ccRNA: CACGGCCGCGCTCGGCTCGA), and a scramble ccRNA (pLV[CRISPR]-hCas9:T2A:Puro-U6>Scramble[gRNA#1], ccRNA: GTGTAGTTCGACCATTCGTG). After lentiviral transduction, cells underwent puromycin selection (4μg/mL) and were single-cell plated into 96 well plates, and allowed to grow for at least 2 weeks. Resulting FASN-deficient clones were screened by western blot.

### Extracellular Flux Analysis

Cellular respiration (oxygen consumption rate (OCR) and extracellular acidification (ECAR)) were assessed using the XF96 Extracellular Analyzer as described in manufacturer’s instructions for the XF cell mito stress and the XF cell mito fuel flex test kits (Agilent Technologies, CA).Briefly, 1.5×10^6^ GL261 or CT2A cells were plated in a 10-cm dish on day 0. Fourty eight hours after plating, cells received a radiation dose of 0 Gy or 10 Gy using the Small Animal Radiation Research Platform (SARRP, Xstrahl Inc, Suwanee GA; open field, 2.82 Gy/min). Immediately after irradiation, 4×10^4^ cells/well were seeded in the XF Cell Culture Microplate and incubated for 24 hours at 37°C. In some conditions cells were incubated with 10 μM of FASN inhibitor (Boehringer Ingelheim; BI-99179). The following day, cells were incubated with 180 μl of Seahorse XF DMEM supplemented with 2 mM L-glutamine, 1 mM sodium pyruvate, and 10 mM glucose for 1 hour prior to assay. The XF Mito Stress Test kit was used to measure mitochondrial respiration as described by the manufacturer. The OCR was measured with sequential injection of 1 μM oligomycin A, 1 μM carbonilcyanide *p*-triflouromethoxyphenylhydrazone (FCCP), and 0.5 μM rotenone/antimycinA. The Seahorse XF Mito Fuel Test kit was also used to measure the mitochondrial fuel usage in live cells. The OCR was measured after injection of 3 μM BPTES (glutamine oxidation inhibitor), 4 μM etomoxir (fatty acid oxidation inhibitor), or 2 μM UK5099 (glycolysis inhibitor). Each inhibitor was injected in a different order to determine the cell dependence on each fuel oxidation pathway. After retrieving the plate from Seahorse instrument, live cell nuclei were stained with NucRed™ Live 647 ReadyProbes™ Reagent (Thermo Fisher Scientific) as per the manufacturer’s instruction. Whole well images were obtained by Spectramax i3x and live cell nuclei were counted (Molecular Devices Jose, CA). All measurements were normalized to the number of live cells in each well and were performed in two biological replicates with 6-12 technical replicates per group.

### Targeted Profiling of Polar Metabolites

1.5×10^6^ GL261 cells were plated in a 10-cm dish on day 0. Twenty four hours after incubation, cells received a radiation dose escalation from 0 Gy, 6 Gy and 10 Gy (n=3 per condition) using the SARRP (open field, 2.82 Gy/min). Immediately after irradiation, medium was replaced in each dish, including the control. Twenty four hours after irradiation, cells were washed twice with cold PBS.The liquid was removed from the dish as thoroughly as possible. The dish was then placed on dry ice. One mL of cold 80% methanol was added to the 10-cm dish prior to cell scraping. Lysates were then transferred into a cold microcentrifuge tube, vortexed 1 min, left on dry ice for 5 min and vortexed 1 min again. Next, lysates were centrifuged at 14,000 x g for 20 min at 4°C. Lysates were then transferred to a 1.5mL cold microcentrifuge tube and stored at -80°C until mass spectrometry analysis at the Weill Cornell Medicine Proteomics and Metabolomics Core Facility. Polar metabolite targeted profiling was performed according to a method described in a previous publication (17). Metabolites were extracted from cells using pre-chilled 80% methanol (-80 °C). The extract was dried completely with a Speedvac (no heat). The dried sample was redissolved in HPLC grade water before it was analyzed with hydrophilic interaction liquid chromatography (HILC) mass spectrometry (MS). Metabolites were measured on a Vanquish UPLC system (ThemoScientific) via an Ion Max ion source with HESI II probe (Thermo Scientific), coupled to a Q Exactive Orbitrap mass spectrometer (Thermo Scientific). A Sequant ZIC-pHILIC column (2.1 mm i.d. × 150 mm, particle size of 5 μm, Millipore Sigma) was used for separation of metabolites. A 2.1 × 20 mm heated guard column with the same packing material was used for protection of the analytical column. Flow rate was set at 150 μL/min. Buffers consisted of 100% acetonitrile for mobile phase A, and 0.1% NH_4_OH/20 mM CH_3_COONH_4_ in water for mobile phase B. The chromatographic gradient ran from 85% to 30% A in 20 min followed by a wash with 30% A and re-equilibration at 85% A. The Q Exactive was operated in full scan, polarity-switching mode with the following parameters: the spray voltage 3.0 kV, the heated capillary temperature 300 °C, the HESI probe temperature 350 °C, the sheath gas flow 40 units, the auxiliary gas flow 15 units. MS data acquisition was performed in the *m/z* range of 70–1,000, with 70,000 resolution (at 200 *m/z*). The AGC target was 1e6 and the maximum injection time was 250 ms. The MS data was processed using XCalibur 4.1 (Thermo Scientific) to obtain signal intensity of each metabolite for relative quantification. Metabolites were identified using an in-house library established using chemical standards. Identification required accurate mass (within 5 ppm) and standard retention times. The intensity values were normalized to total cell count prior downstream analysis.

### Free Fatty Acid Profiling

1.5×10^6^ GL261 or CT2A cells were plated in a 10-cm dish on day 0. Twenty four hours after plating, cells received a radiation dose escalation from 0 Gy, 6 Gy and 10 Gy (n=5 per condition) using the SARRP (open field, 2.82 Gy/min). Immediately after irradiation, medium was replaced in each dish, including the control. Twenty four hours after irradiation, cells were detached from plates with trypsin, washed twice with cold PBS and spun down for 5 min at 340 x g. Cells were then transferred to a 1.5mL centrifuge tube and stored at -80°C until mass spectrometry analysis at the Weill Cornell Medicine Proteomics and Metabolomics Core Facility.

Briefly, free fatty acids were extracted from cells using 90% methanol (LC-MS grade, Thermo Scientific). The extracts were clarified by centrifugation at 15,000 x g for 10 min. The supernatant was collected and dried down using a SpeedVac. The dried sample was reconstituted using 50% methanol prior to LC-MS analysis.

Chromatographic separation was performed on a Vanquish UHPLC system (Thermo Scientific) with a Cadenza CD-C18 3 μm packing column (Imtakt, 2.1 mm id x 150 mm) coupled to a Q Exactive Orbitrap mass spectrometer (Thermo Scientific) via an Ion Max ion source with a HESI II probe (Thermo Scientific). The mobile phase consisted of buffer A: 5 mM ammonium acetate in water (LC-MS grade, Thermo Scientific) and buffer B: 5 mM ammonium acetate, 85% isopropanol (LC-MS grade, Thermo Scientific), 10% acetonitrile (LC-MS grade, Thermo Scientific), and 5% water. The LC gradient was as follows: 0–1.5 min, 50% buffer B; 1.5-3 min, 50-90% buffer B; 3-5.5 min, 90-95% buffer B; 5.5-10 min, 95% buffer B, followed by 5 min of re-equilibration of the column before the next run. The flow rate was 150 μL/min. MS data was acquired in negative ionization mode. The following electrospray parameters were used: spray voltage 3.0 kV, heated capillary temperature 350 °C, HESI probe temperature 350 °C,sheath gas, 35 units; auxiliary gas 10 units. For MS scans: mass scan range, 140-1000 *m/z*; resolution, 70,000 (at *m/z* 200); automatic gain control target, 1e6; maximum injection time, 50 ms.

MS data files were processed using XCalibur (version 4.1, Thermo Scientific). Identification of free fatty acids was based on accurate masses within 5 ppm and standard retention times from compound standards. Relative quantitation was performed based on MS signal intensities. The intensity values were normalized by the total protein in each sample (determined by the Pierce™ BCA Protein assay kit, ThermoFisher Scientific, cat# 23225) before they were used for quantitative analysis.

### Targeted Profiling of Polar Metabolites and Free Fatty Acid Profiling Analysis

Statistical analysis was performed with the online software MetaboAnalyst 5.0 (18). Analysis of polar metabolites was constricted to metabolites that were common between the two independent experiments of GL261. Data were not transformed, but were Pareto scaled for both sets of analyses. Statistical analysis was performed with an ANOVA with an adjusted p-value (false discovery rate FDR) cutoff of <0.05 and Tukey’s as the post-hoc analysis. Fatty acid analysis was based on the comparison between GL261 and CT2A on the statistically significant metabolites. Data were visualized in the form of a principal component analysis (PCA) scores plot based on the first two components and in heatmaps with Euclidean distance measure and Ward clustering. Enrichment analysis of the polar metabolites was KEGG pathway based and only metabolite sets that contained at least 5 entries were used.

### Tumor Challenge and Treatment

On day 0, tumor cells (i.e. GL261 or CT2A) for intracerebral implantation were trypsinized, counted and checked for viability by trypan blue exclusion. Cells were washed twice with DMEM without FBS or antibiotics, and a final suspension of 25×10^6^ cells/mL in DMEM was prepared. Mice were anesthesized with 1.5% isoflurane. Using a stereotactic head frame and a 5μL Hamilton syringe (Hamilton glass syringe 700 series RN), 5×10^4^ of CT2A cells in 2μL were injected into the mice right striatum. The coordinates used for intracerebral injection were 0.5mm anterior to the bregma, 2mm lateral to the saggital suture (right hemisphere), and 3 mm below the dura. Mice with established CT2A tumors were selectively irradiated with 0Gy or 10Gy ten days after tumor cells implantation as previously described (19). Briefly, mice were anesthetized with isoflurane to perform computed tomographic (CT) imaging on the SARRP to identify the burr hole from tumor implantation, which was used to aim 10Gy-radiation in a 3mm beam centered on the tumor. The dose rate was 2.39Gy/min. Criteria to euthanize mice during survival experiment consisted in weight loss of 20% of initial weight, dyspnea, abnormal posture, difficulty with ambulation or any other clinical sign of large intracranial tumor causing significant pain or distress such as lethargy, poor grooming, dehydration, head tilt or hydrocephaly.

### Osmotic Pump Implantation and Removal in Mice

Osmotic pumps were purchased from ALZET (Model 1002) with a manufacture pump rate of 0.25 μL per hour over 14 days. Osmotic pumps were filled with 10 mg/mL of FASN inhibitor (Boehringer Ingelheim; BI-99179) dissolved in 10% DMSO 40% PEG300 50% PBS according to manufacturer’s instructions. The filled pumps were primed by incubation in 1X PBS overnight at 37°C. On the day of implantation (i.e. 10 days after tumor cells implantation), mice were anesthesized with 1.5% isoflurane. After positioning the mice onto a stereotactic frame, a 1-cm sagittal incision was made in the head to expose the skull and a subcutaneous pocket was generated to place the pump. Next, the cannula was positioned using the same stereotactic coordinates of tumor cells injection (i.e. 0.5mm anterior to the bregma, 2mm lateral to right hemisphere, and 3 mm below the dura). A thin layer of adhesive was added to the base of the canula pedestral. Once firmly set, the wound was closed with sutures. Fourteen days post implantation, the osmotic pump was removed.

### Matrix-Assisted Laser Desorption Ionization Imaging Mass Spectrometry (MALDI-IMS)

Brains from tumor-bearing animals were frozen at day 15 (i.e. 5 days after irradiation) in isopentane cooled with dry ice and stored at -80°C. Ten micron cryosections from 3 brains per group were mounted on conductive slides coated with Indium Tin Oxide (ITO) (Delta Technologies; cat # CB-90IN-S111) to perform MALDI-IMS using the Bruker SciMax in an ultrahigh mass resolution Fourier Transform – Ion Cyclotron Resonance (FT-ICR) mass spectrometer. Briefly, ITO-slides with tissue sections are transferred from -80°C to a vacuum chamber and dried for 30 min prior to matrix deposition using N-(1-naphthyl) ethylenediamine dihydrochloride (NEDC, 10mg ml^-1^ in 75% methanol) for negative ion detection and 2,5-Dihydroxybenzoic acid (DHB, 30mg/ml in 90% acetonitrile) for positive ion detection, using a matrix sprayer HTX TM-Sprayer™ (HTX Technologies LLC, NC). Matrix-coated tissue sections are dried in vacuum desiccator for 20 min before loading for MALDI-IMS data acquisition on a 7T Scimax-MRMS mass spectrometer (Bruker Daltonics, USA) equipped with a SmartBeam II laser and a MALDI source (spatial resolution set at 80μm). Acquired image data were exported to Scils Lab software (SCiLS, Bremen, Germany) and Cardinal R package (https://bioconductor.org/packages/release/bioc/html/Cardinal.html) for image visualization and statistical analysis. Compound identifications are assigned based on both accurate mass and isotope pattern matches to HMDB (Human Metabolome Database) (<3 ppm). Mean image intensities from corresponding regions were compared using Student t-tests (2-tailed).

### Hematoxylin and Eosin Staining of brain tissue sections

Ten micron cryosections from tumor-bearing brains were stained with Hematoxylin and Eosin (H&E) according to the manufacturer’s instruction (Abcam; cat# ab245880). Briefly, sections were fixed with 4% PFA in PBS for 10 minutes then washed twice in PBS. Slides were incubated with Hematoxylin, Mayer’s (Lillie’s Modification) for 2 minutes at room temperature, then rinsed twice in distilled water to remove the excess of stain. Next, slides were incubated with Bluing Reagent for 10-15 seconds, rinsed twice with distilled water and dip in absolute alcohol. After blotting the excess off the slide, Eosin Y Solution (Modified Alcoholic) was applied to cover the brain sections and incubated for 2 minutes. Slides were then rinsed with absolute alcohol, dehydrated with three changes of absolute alcohol, cleared in xylene and mounted for analysis.

### Evaluation of intracellular neutral lipids

To determine the content of neutral lipids in irradiated glioblastoma cells, 1×10^6^ GL261 or CT2A were plated in a 10-cm tissue culture dish on day 0. Twenty four hours after incubation, cells received a radiation dose escalation from 0 Gy to 10 Gy using the SARRP (open field, 2.82 Gy/min). Immediately after radiation, medium was replaced from each dish, including the control. In some conditions cells were incubated with 10 μM of FASN inhibitors BI-99179 (Boehringer Ingelheim) or TVB-3166 (Selleck Chemicals). Twenty four hours, 2 days, 3 days, 6 days or 7 days after irradiation, cells were collected and stained for neutral lipids by BODIPY™ 493/503 (4,4-Difluoro-1,3,5,7,8-Pentamethyl-4-Bora-3a, 4a-Diaza-s-Indacene; ThermoFisher Scientific; cat# D3922) or HCS LipidTOX™ Red Neutral Lipid Stain (ThermoFisher Scientific; cat# H34476) according to the manufacturer’s intructions. Briefly, cells were trypsinized, washed with PBS, and incubated with 10μM of BODIPY or 1:200 LipidTOX staining solution in PBS in the dark for 15 min at 37°C. After incubation, cells were washed with PBS. Cells were then centrifuged for 5 min at 340 x g and resuspended with 200μL of 4% PFA in PBS (Santa Cruz Biotech, cat# sc-281692). All samples were acquired with a MACSQuant Analyzer 10 flow cytometer and analyzed with FloJo software version 10.4.0 (Tree Star). Results are displayed as mean fluorescence intensity with n=3 per group. Experiments were done at least in two biological experiments.

For LipidTOX staining for microscopy, 1.3×10^4^ GL261 or 1.8×10^4^CT2A cells were plated in 24-well plates with 12 mm glass round coverslips on day 0. Twenty four hours after incubation, cells were exposed to 0 Gy or 10 Gy using the SARRP (open field, 2.82 Gy/min). Immediately after irradiation, medium was replaced in each dish, including the control. Twenty four hours after irradiation, cells were washed with PBS, and incubated with 1:200 LipidTOX staining solution in PBS in the dark for 15 min at 37°C. After incubation, cells were washed with PBS and then fixed with 4% PFA in PBS for 30 min at room temperature. Coverslips were mounted onto glass slides using a mounting medium containing DAPI (Vector Laboratories, cat# H-1800). Images were obtained using a Zeiss Axioplan Upright Wide Field Microscope at the Weill Cornell Medicine Optical Microscopy Core Facility.

### Quantification of Prostaglandin E2

1×10^6^ GL261 or CT2A cells were plated in 10-cm dishes on day 0. Twenty four hours after incubation, cells received a radiation dose of 0 Gy or 10 Gy (SARRP; open field, 2.82 Gy/min). Immediately after irradiation, medium was replaced in each dish, including the control. Supernatants were collected 24h after completion of tumor cells irradiation, and live cells were counted using Trypan Blue. Next, prostaglandin E2 (PGE2) was measured in cell-free supernantants using a sequential competitive enzyme immunoassay (Prostaglandin E2 Parameter Assay Kit, cat# KGE004B, R&D Systems) following the manufacturer’s instructions. Results were normalized by the number of live cells.

### Immunostaining of GBM cells

1.3×10^3^ GL261 or 1.8×10^3^ CT2A cells were plated in 24-well plates with 12mm glass round coverslips on day 0. Twenty four hours after incubation, cells were exposed to 0 Gy, 6 Gy and 10 Gy using the SARRP (open field, 2.82 Gy/min). Immediately after irradiation, medium was replaced in each dish, including the control. Twenty four hours after irradiation, cells were fixed with 4% PFA in PBS (Santa Cruz Biotech, cat# sc-281692). Cells were permeabilized and blocked simultaneously with 1% BSA (Gemini Bio-Products, cat# 700-100P), 10% normal goat serum (Cell Signaling Technology, cat# 5425S), 0.3M glycine (Bio-Rad, cat# 161-0717), 0.1%Triton X-100 (Fisher, cat# PB151) in PBS for 1 hour at room temperature. Cells were incubated with a primary anti-FASN antibody diluted in 1%BSA PBS-0.05%Tween 20 (Abcam, cat# ab22759; 1:200) or Normal Rabbit IgG for the isotype control (Cell Signaling, cat# 2729; 1:200) overnight at 4°C. After three washes in PBS-0.05%Tween 20, samples were incubated with a secondary antibody diluted in 1%BSA PBS-0.05%Tween 20 (goat anti-mouse IgG H&L – Alexa fluor 488 preabsorbed, Abcam, cat# ab150081; 1:500) for 1 hour at room temperature. After 2 washes in PBS-0.05%Tween 20 then 2 washes in PBS, mounting medium containing DAPI was used (Vector Laboratories, cat# H-1800) to counterstain nuclei. Images were obtained using a Zeiss Axioplan Upright Wide Field Microscope at the Weill Cornell Medicine Optical Microscopy Core Facility . Semi-automated quantification of FASN expression was performed by using Fiji software (v 1.53n; (20)) on at least 10 images for each condition. Integrated density was measured for all the cells of each images using a threshold to exclude the background. Then, the sum of the integrated density of an image was divided by the number of nuclei counted in that image in order to obtain the mean integrated density of cell per image.

### Clonogenic assay

0.8×10^6^ GL261 or CT2A cells were plated in 10-cm dishes on day 0. On day 2, cells received a radiation of 0 Gy or 4 Gy using the SARRP irradiator (open field, 2.82 Gy/min). Immediately after irradiation, medium was replaced in each dish, including the control. Three hours after irradiation, cells were detached from plates with trypsin, resuspended in complete media and counted. Next, cells were plated into 6-well plates at the density of 2.5×10^2^, 5×10^2^, 10×10^2^ for GL261 and 1×10^2^, 2×10^2^, 4×10^2^ for CT2A. FASN inhibitor (Boehringer Ingelheim; BI-99179) was added to the wells at a concentration of 1μM. After 8-12 days, the cells were fixed with 70% ethanol for 10 minutes and washed with distilled water. Cells were then stained with 0.1% crystal violet (Sigma-Aldrich, #V5265) for 15 minutes, washed with distilled water and air dried overnight. Plates were imaged for red fluorescence (crystal violet has absorption/emission peaks at 595/635 nm, respectively) on a C600 Gel Doc and Western Imaging System (Azure Biosystems). The colonies consisting of at least 50 cells were counted in each well by a semi-automatic method using Fiji software (20). Survival fractions were calculated as previously described (21).

### Flow Cytometry Analysis of Apoptotic and Necrotic Cells

To determine the induction of apoptosis and necrosis in irradiated GBM cells, 1×10^6^ GL261 or CT2A derivatives (i.e. GBMsgControl and GBMsgFasn) were plated in a 10 cm tissue culture dish on day 0. Twenty four hours after incubation, cells received a radiation dose of 0 Gy or 10 Gy (n=3 per condition; SARRP; open field, 2.82 Gy/min). Immediately after irradiation, culture medium was replaced in each dish, including the control. In some conditions cells were incubated with 10 μM of FASN inhibitor (Boehringer Ingelheim; BI-99179). Twenty four hours after irradiation cells were trypsinized and stained using the FITC Annexin V Apoptosis Detection Kit with PI (Biolegend, CA) following the manufacturer’s instructions. All samples were acquired with a MACSQuant Analyzer 10 flow cytometer and analyzed with FloJo software version 10.4.0 (Tree Star). Cell populations were separated as follows: Apoptotic cells: Annexin V+/PI-, Necrotic cells: Annexin V+/PI+ and expressed as % of total single cells.

### Transmission Electronic microscopy

GBM cells were incubated with 4μg/mL of doxycycline 10 days prior to treatment to induce the knockdown of *Fasn*. Next GL261 or CT2A derivatives were irradiated with a radiation dose of 0 Gy or 10 Gy. Twenty four hours after irradiation, cells were washed and fixed for 30 minutes in 2.5% glutaraldehyde, 4% paraformaldehyde and 0.002% of picric acid. Next, cells were washed with PBS and post-fixed with osmium (2% osminumtetroxide, 3 K-ferricyanide, 1:1 aqu) for 30-60 minutes at room temperature. The cells were then washed and stained en-bloc with 1.5% uranul acetate (aqu) for 30 minutes followed by dehydration in graded ethanol series 50%, 70%, 85%. 95%, 3×100% (5-10 min at each step). The cells were finally embedded in Epon-analog resin comprised of 9.7 g of LX112 (Ladd Research Industries), 3.2g of DDSA (Electron microscopy sciences), 5.9 g of NMA (Electron microscopy sciences) and 34mL/mL of BDMA (Elextron microscopy sciences). Sections (1μm) were produced on a EM UC6 ultramicrotome and stained with 2% uranyl acetate and Reynold’s lead citrate. Transmission electronic microscopy (TEM) was performed on JEM-1400 transmission electron microscope at 80 kV. These experiments were perfomed at the Weill Cornell Medicine Microscopy Core Facility.

### Quantification of HMGB1

GBM cells were incubated with 4μg/mL of doxycycline 10 days prior to treatment to induce the knockdown of *Fasn*. Next, 0.5×10^5^ cells of GL261 or CT2A derivatives (i.e. GL261shNS and GL261shFASN) were plated in a 10-cm dish on day 0. Twenty four hours after incubation, cells received a radiation dose of 0 Gy or 10 Gy (n=3 per condition; SARRP; open field, 2.82 Gy/min). Immediately after irradiation, medium containing doxycycline was replaced in each dish, including the control. Twenty four hours after completion of tumor cells irradiation, supernatants were collected and live cells were counted using Trypan Blue. Next, HMGB1 was measured in cell-free supernantants using an enzyme-linked immunosorbent assay (Tecan HMGB1 ELISA, cat# ST51011, Fisher Scientific) following the manufacturer’s instructions. Results were normalized by the number of live cells.

### Western Blot

Proteins from GL261 and CT2A cells were extracted in RIPA buffer (Sigma, cat# R0278-50mL) containing a protease inhibitor cocktail (Sigma, cat # 11836153001) and a phosphatase inhibitors cocktail (ThermoFisher Scientific, cat # 78420). Protein concentrations were determine with BCA method (ThermoFisher Scientific, cat # 23227). A total of 30-50μg of protein of each sample was loaded on a 8-10% SDS-PAGE and run at 100V constant voltage. Transfer of protein on activated PVDF membranes (EMD-Millipore, cat#IPVH07850) were performed overnight at 4°C at 50mA. PVDF membraned were probed with primary antibodies (m-h-anti-BIP 1:1000, cat #3177T; m-h-anti-CHOP 1:1000, m-h-anti-fatty acid synthase 1:100, cat#3180T; m-h-anti-vinculin 1:1000, cat#13901S; Cell Signaling Technology) at 4°C overnight. Blots were incubated with ECL anti-rabbit IgG secondary antibody (Sigma, cat# GENA934-1 mL; 1:5000) at room temperature for 1 hour. Chemiluminescence was used to visualize protein bands (Perkin Elmer, cat# NEL103001EA). Pictures were acquired using the Azure c600 imager (Azure Biosystems). Precision Plus Protein™ Western C™ Blotting standards and Precision Protein™ Streptactin-HRP conjugate were used to estimate molecular weight (BioRad, cat# 1610380 and cat#1610376). Relative quantification was performed using ImageJ software (version 1.53p (22)) to determine the protein expression ratios for each sample.

### Processing and analysis of human glioblastoma data

GBM expression data were downloaded from the Glioma Longitudinal Analysis (GLASS) Consortium as normalized transcripts per million (TPM) (23). Patients were filtered using the following characterizations: Glioblastoma histology, ≥50Gy radiation dose received between primary tumor (TP) and first recurrence (R1), and paired TPM RNA-sequencing (RNA-seq) counts available at TP and R1 time points. After filtering, we analyzed 55 paired TP-R1 samples.

We next performed gene set variation analysis (GSVA) (24) on pathways of interest. A list of relevant pathways was constructed using the C2 (includes canonical pathways from bonafide sources such as Biocarta, Kegg, PID, Reactome, and Wikipathways), C5 (gene and phenotype ontology pathways), and H (hallmark gene sets) human geneset collections from the Molecular Signatures Database (MSigDB) (25). Pathways of interest were extracted using keywords “LIPID” and “FATTY_ACID” excluding 24 “GARGALOVIC_” pathways due to irrelevance. Ultimately, 193 genesets meeting the additional requirement of containing between 10 and 500 genes were run using GSVA on each TP and R1 expression counts from the 55 GBM patients specified above. Using the scores generated by GSVA, a paired t-test was run to assess the significance of the score change between TP and R1 for each pathway. Additionally, a false discovery rate (FDR) test was run on the p-values. P-value significance was determined by p < 0.05 (n = 46/193), and FDR significance was determined by an FDR < 0.05 (n = 1/193). For ridge plot visualization, only pathways significant by p-value are shown with their distribution of delta values (score at R1 – score TP). An overall increase of pathway score at recurrence is determined by an average delta score of > 0.

We next evaluated differential gene expression of recurrent and primary human glioblastoma tumors using REACTOME_METABOLISM_OF_LIPIDS pathway downloaded from the MSigDB. This pathway was excluded from the previous GSVA analysis as it contains more than 500 genes. However, it was deemed most appropriate for differential gene expression analysis as it compiles a broad list of genes relevant to lipid metabolism. The TPM files were subsetted to only include expression reads from the genes on this list (n = 712). Differential gene expression analysis comparing paired TP and R1 human glioblastoma tumors was then conducted using DESeq2 (26). For volcano plot visualization, only genes with an adjusted p-value < 0.05 (by Benjamini and Hochberg’s correction for multiple comparisons) and log2 fold change of > |0.8| were labelled as significant. For the “REACTOME_FATTY_ACIDS” pathway, individual GSVA scores for each matched TP and R1 case were plotted and a line was added between the two to visualize patientspecific pathway score changes between time points. A Wilcoxon signed-rank test was used to evaluate significance between the positive (n = 39) and negative deltas (n = 16).

### Statistics

All statistical analyses were performed using GraphPad Prism 9 and are considered significant when *p*<0.05. The identity of the statistical test performed, *p* values and n values are stated in the figure legends. One-way Analysis of Variance (ANOVA) followed by Tukey’s post hoc multiple comparisons was applied for statistical analysis of neutral lipids accumulation, free fatty acid profiles, HMGB1 quantification, TEM analysis of mitochondria, targeted metabolic profiles, FASN expression by immunocytochemistry, flow cytometry analysis of apoptotic and necrotic cells and metabolic fuel flux. Unpaired student’s t test was applied for statistical analysis of Seahorse MitoStress Assay, MALDI-IMS, PGE2 quantification and clonogenic assay. Log-rank (Mantel-Cox) test was used for statistical anlysis of Kaplan Meyer survival curves. Wilcoxon signed-rank tests was used for comparisons of primary and recurrent tumors (Fig. 6C) since pairs of samples from the same patient had to be compared and a normal distribution was not always given.

**Figure 1:**
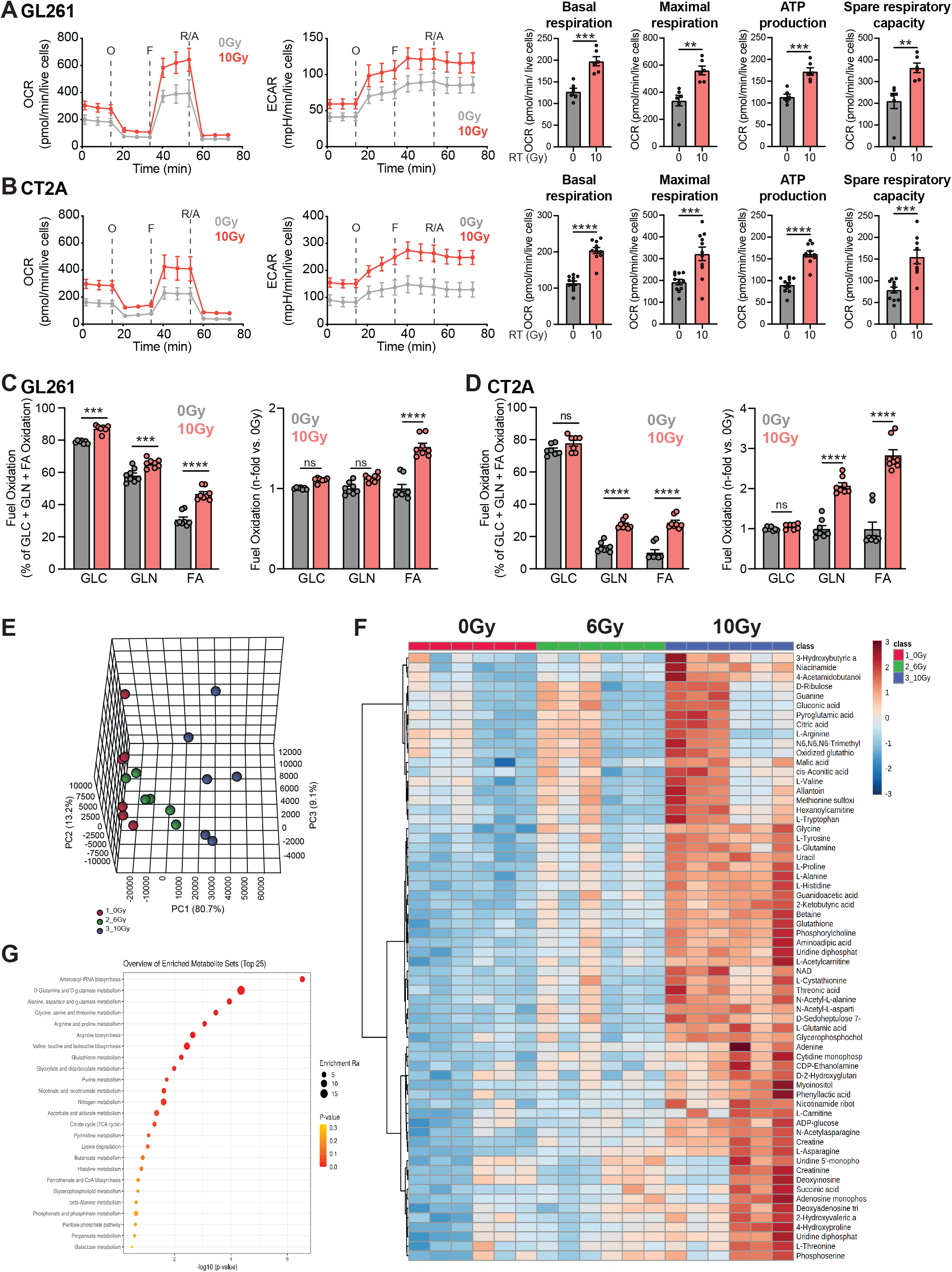
Radiation induces metabolic reprogramming in glioblastoma cells. Murine glioblastoma cells GL261 and CT2A were analyzed 24hrs after irradiation (10 Gy) by extracellular flux analysis. Fundamental mitochondrial function was determined by Seahorse MitoStress Assay for oxygen consumption rate (OCR) and extracellular acidification rate (ECAR) at baseline and in response to sequential injections of oligomycin (O), FCCP (F), and rotenone with antimycin (R/A) in GL261 (**A**) and CT2A (**B**). OCR was used to quantify ATP production and spare respiratory capacity (SRC). Substrate fuel capacity for the three major fuel pathways of glucose (GLC), glutamine (GLN) and fatty acid (FA) was determined by Seahorse Mito Fuel Flex Test in GL261 (**C**) and CT2A (**D**). Values represent mean ± SEM. ns, not significant; **p<0.01; ***p<0.001; ****p<0.0001, using unpaired Student’s t test (**A** and **B**), or one-way ANOVA followed by Tukey post hoc comparisons (**C** and **D**). (**A – D**) Experiment was done in duplicate with n=6-12 per group. (**E – G**) GL261 cells were analyzed by targeted metabolic profiling 24hrs after irradiation with 6 Gy or 10 Gy. Metabolites commonly detected in two independent (n=3/group) experiments were combined for analysis. (**E**) PCA score plots. (**F**) Heatmap of differentially represented metabolites in 6Gy and 10Gy. (**G**) Pathway enrichment analysis in common metabolites between two independent experiments. Experiments were done in duplicate with n=3 per group.

**Figure 2:**
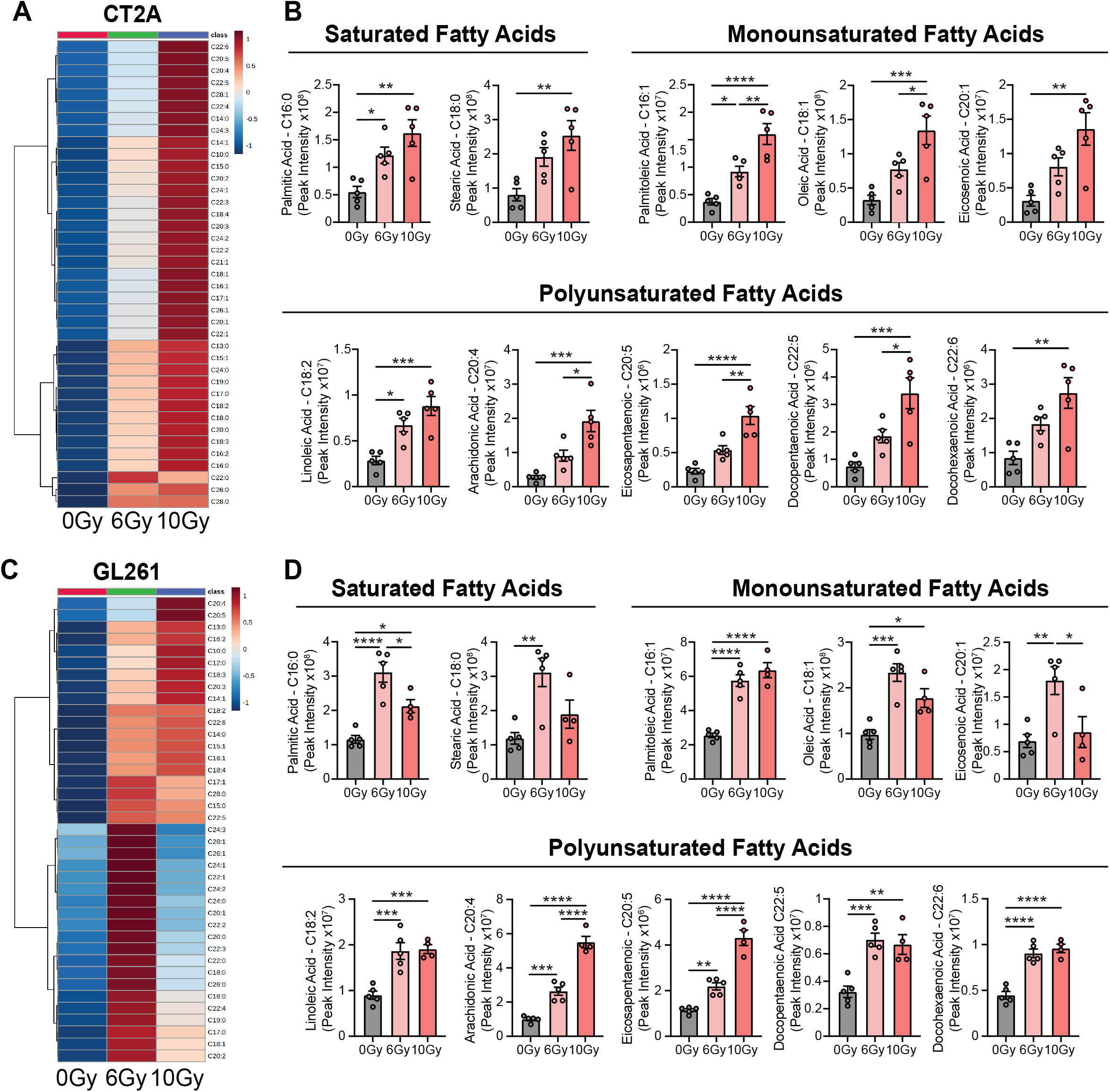
Unsaturated fatty acids are preferentially induced by irradiation of glioblastoma cells *in vitro*. GL261 and CT2A cells were subjected to 6 Gy or 10 Gy irradiation *in vitro* and 24hrs later cells were collected for free fatty acid profiling by mass spectrometry. Heatmap of differentially represented fatty acids in averaged replicates (n=5/group) in CT2A (**A**) and GL261 (**C**) cells. Peak intensity of representative saturated, monounsaturated, and polyunsaturated fatty acids for CT2A (**B**) and GL261 (**D**) cells. Values represent mean ± SEM. Each dot represents one replicate. *p<0.05; **p<0.01; ***p<0.001; ****p<0.0001, using one-way ANOVA followed by Tukey post hoc comparisons (B and D). Experiment was done in duplicate with n=5 per group.

**Figure 3:**
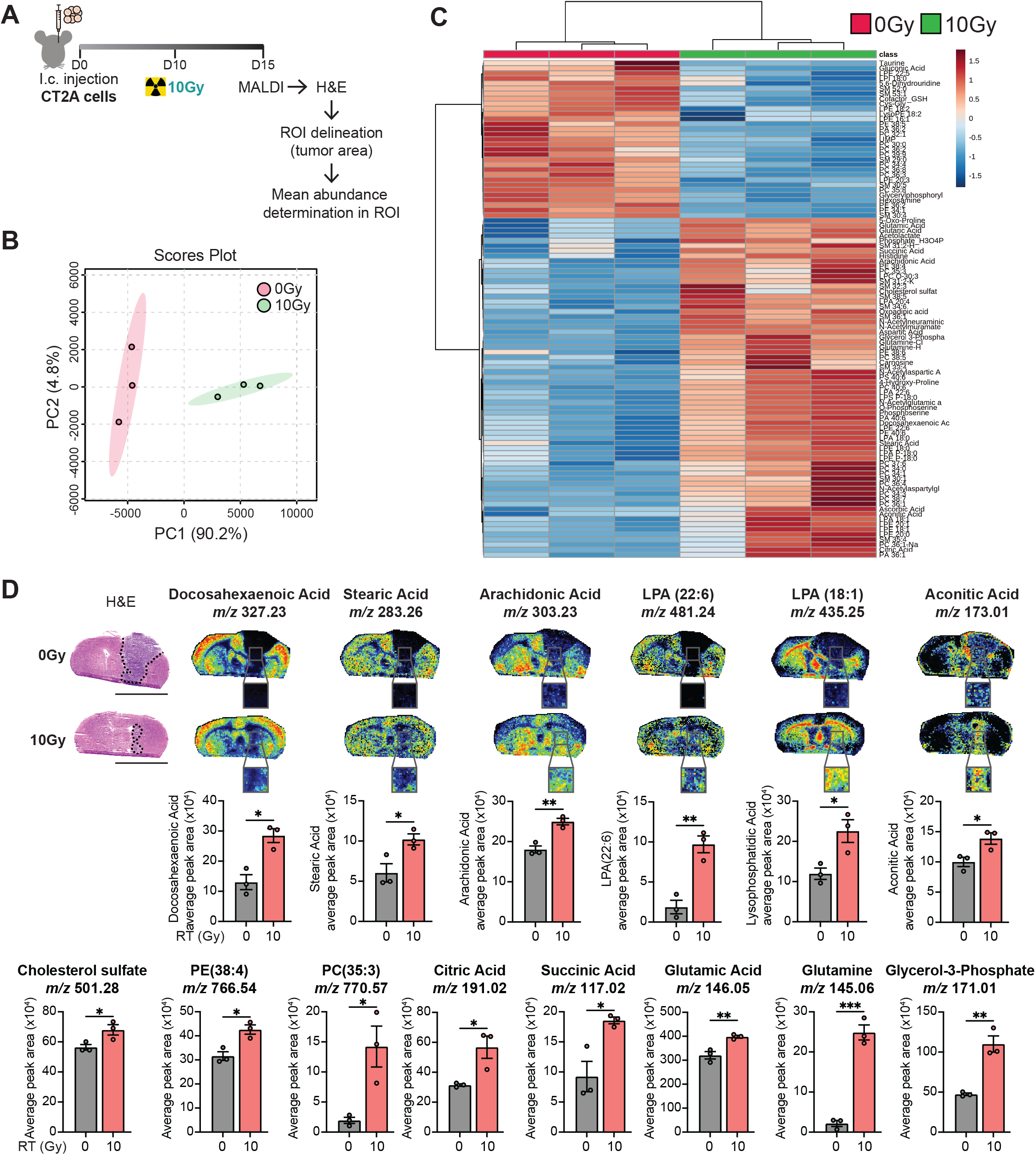
Radiation alters glioblastoma metabolism toward the generation of unsaturated fatty acids *in vivo*. (**A**) Experiment scheme and timeline. Briefly, 50,000 CT2A cells were injected i.c. into C57BL/6N mice on day 0 (n=3/group). On day 10, radiation was selectively given to i.c. tumors in a single dose of 10 Gy. On day 15, tumor-bearing brains were collected for MALDI imaging. Metabolites were quantified in the tumor area delineated with black doted lines in the H&E sections. (**B**) PCA score plots. (**C**) Heatmap of differentially represented metabolites in 10Gy irradiated tumors. (**D**) H&E of brain sections from animals in (A). Scale 5mm. Peak intensity and representative images of metabolites. Values represent mean ± SEM. Each dot represents one animal. *p<0.05; **p<0.01; ***p<0.001, using unpaired Student’s t test (D).

**Figure 4:**
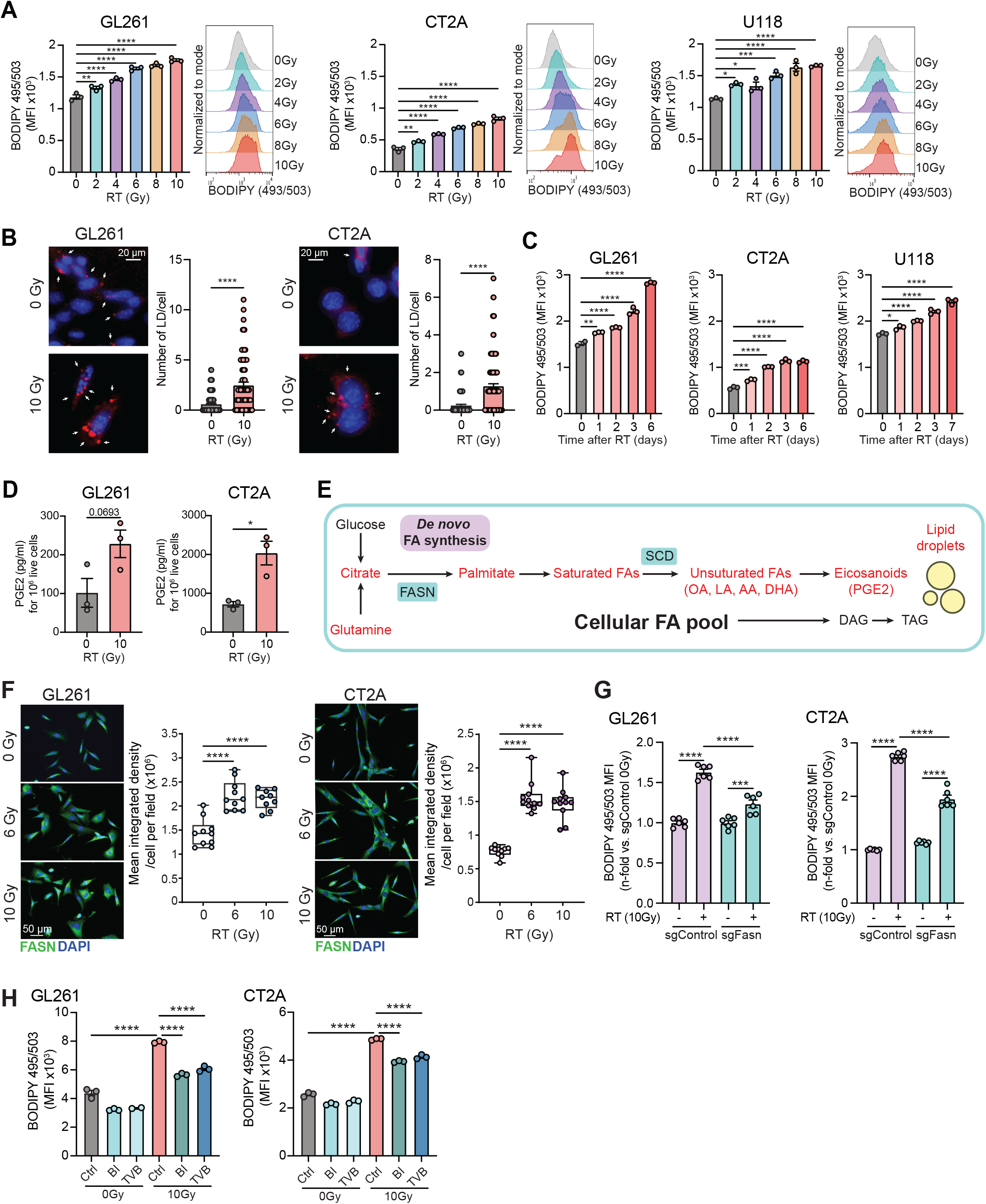
Radiation induces FASN-mediated accumulation of lipids in glioblastoma cells. (**A**) Murine glioblastoma GL261 and CT2A cells and human glioblastoma U118 cells were subjected to different radiation doses from 2 Gy to 10 Gy *in vitro* and 24hrs later, neutral lipids accumulation was determined by BODIPY (495/503) staining and flow cytometry. (**B**) Representative images and quantification of LipidTOX fluorescence staining for 10 Gy irradiated GL261 and CT2A cells. Scale 20μm. Lipid droplets are pointed with white arrows. (**C**) Glioblastoma cells were plated on day 0 and subjected to 10 Gy irradiation at different time points (from 1 to 7 days before collection) to synchronize the collection and BODIPY staining on the same day. (**D**) PGE2 secretion was determined by ELISA 24hrs after 10 Gy irradiation of GL261 and CT2A cells. Results are expressed in pg/mL ± SEM for 1,000,000 live cells. (**E**) Overview of the main pathways responsible for the cellular fatty acid pool: *de novo* fatty acid synthesis and uptake. Highlighted in red are the components described to be modulated by radiation (Figs. 1-4). (**F**) Representative images and quantification of FASN immunofluorescence staining 24hrs after 6 Gy and 10 Gy irradiation in GL261 and CT2A cells. Scale 50μm. (**G**) Flow cytometry analysis of BODIPY 495/503 staining in CRISPR/Cas9 sgControl or sgFasn GL261 and CT2A cells 24hrs after 10 Gy irradiation. (**H**) Flow cytometry analysis of BODIPY 495/503 staining in GL261 and CT2A cells 24hrs after 10Gy irradiation and treatment with the FASN inhibitors 10 μM BI-99179 (BI) and TVB-3166 (TVB). Values represent mean ± SEM. Each dot represents one replicate. *p<0.05; **p<0.01; ***p<0.001; ****p<0.0001, using one-way ANOVA followed by Tukey post hoc comparisons (A and F-H) or unpaired Student’s t test (B and D). Experiments were done in duplicate with n=3 per group.

**Figure 5:**
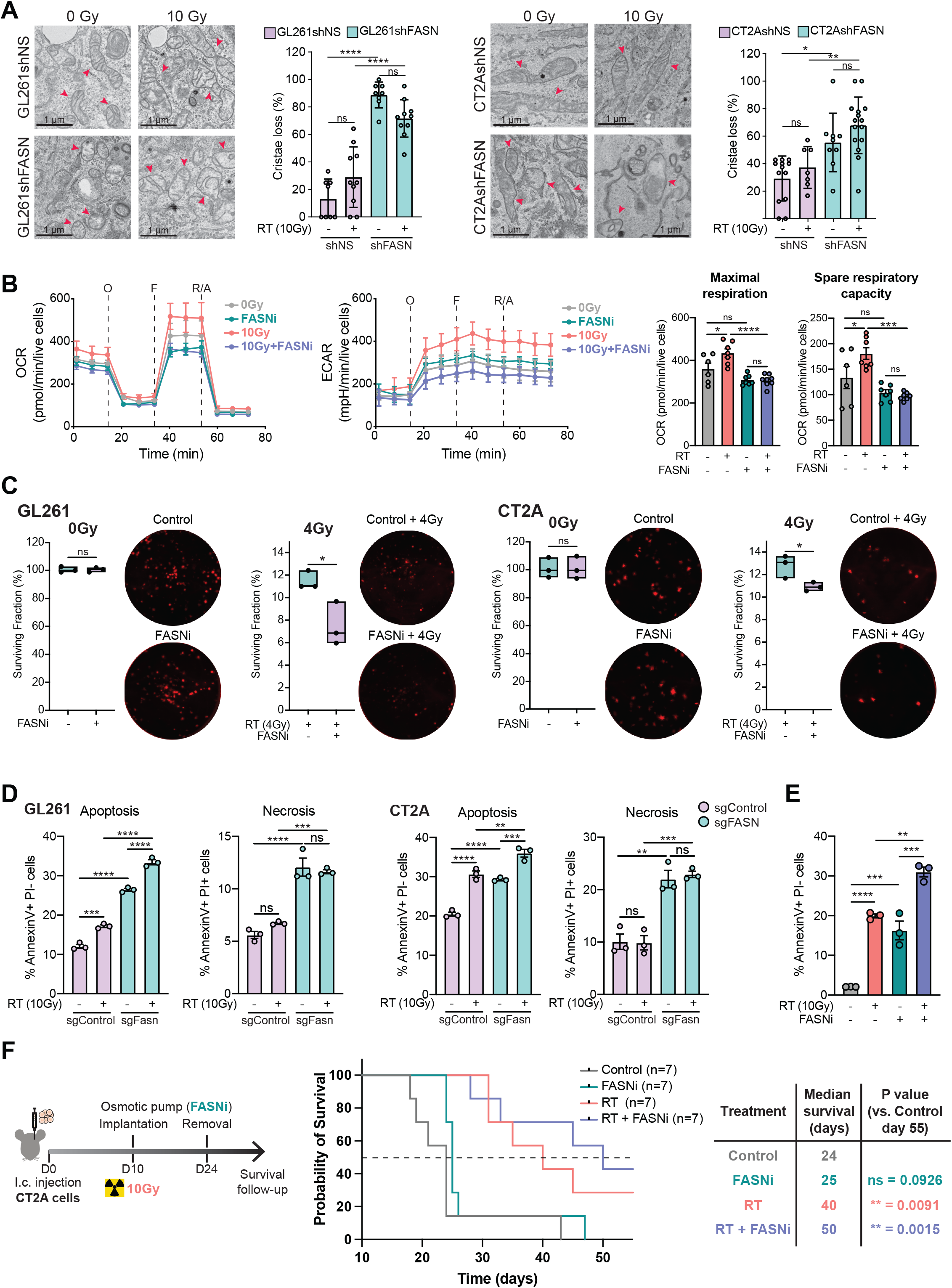
FASN inhibition induces apoptosis and tumor control of irradiated GBM. (**A**) Representative transmission electron microscopy images and quantification of mitochondria without cristae in 10 Gy irradiated shNon-silencing (shNS) or shFASN GL261 and CT2A cells. Scale 1 μm. (**B**) Mitochondrial function was determined by Seahorse MitoStress Assay for oxygen consumption rate (OCR) and extracellular acidification rate (ECAR) at baseline and in response to sequential injections of oligomycin (O), FCCP (F), and rotenone with antimycin (R/A) in CT2A cells treated for 24hrs with 10 Gy and/or 10 μM FASNi (BI-99179). OCR was used to quantify spare respiratory capacity. (**C**) GL261 and CT2A cells that received 4 Gy irradiation were cultured for 2 weeks in the presence 10 μM of the FASN inhibitor (FASNi), BI-99179, fixed, and stained with crystal violet. Representative images of the colonies and quantitative data are reported. Flow cytometry analysis of AnnexinV/PI staining 24hrs after 10 Gy irradiation in CRISPR/Cas9 sgControl or sgFasn GL261 and CT2A cells (**D**) and CT2A cells treated with 10 μM FASNi (BI-99179) (**E**). Values represent mean ± SEM. Each dot represents one replicate. ns, not significant; *p<0.05; **p<0.01; ***p<0.001; ****p<0.0001, using unpaired Student’s t test (A) or one-way ANOVA followed by Tukey post hoc comparisons (B-E). Experiments were done in duplicate with n=3 per group. (**F**) Experiment scheme and timeline. Briefly, 50,000 CT2A cells were injected i.c. into C57BL/6N mice on day 0 (n=7/group). On day 10, radiation was selectively given to i.c. tumors in a single dose of 10 Gy. FASNi (BI-99179) was administered via osmotic pump/brain infusion kit between days 10–24. Survival was monitored for 55 days. Kaplan Meyer survival curves and median survival values are reported. **p<0.01, using Log-rank (Mantel-Cox) test.

**Figure 6:**
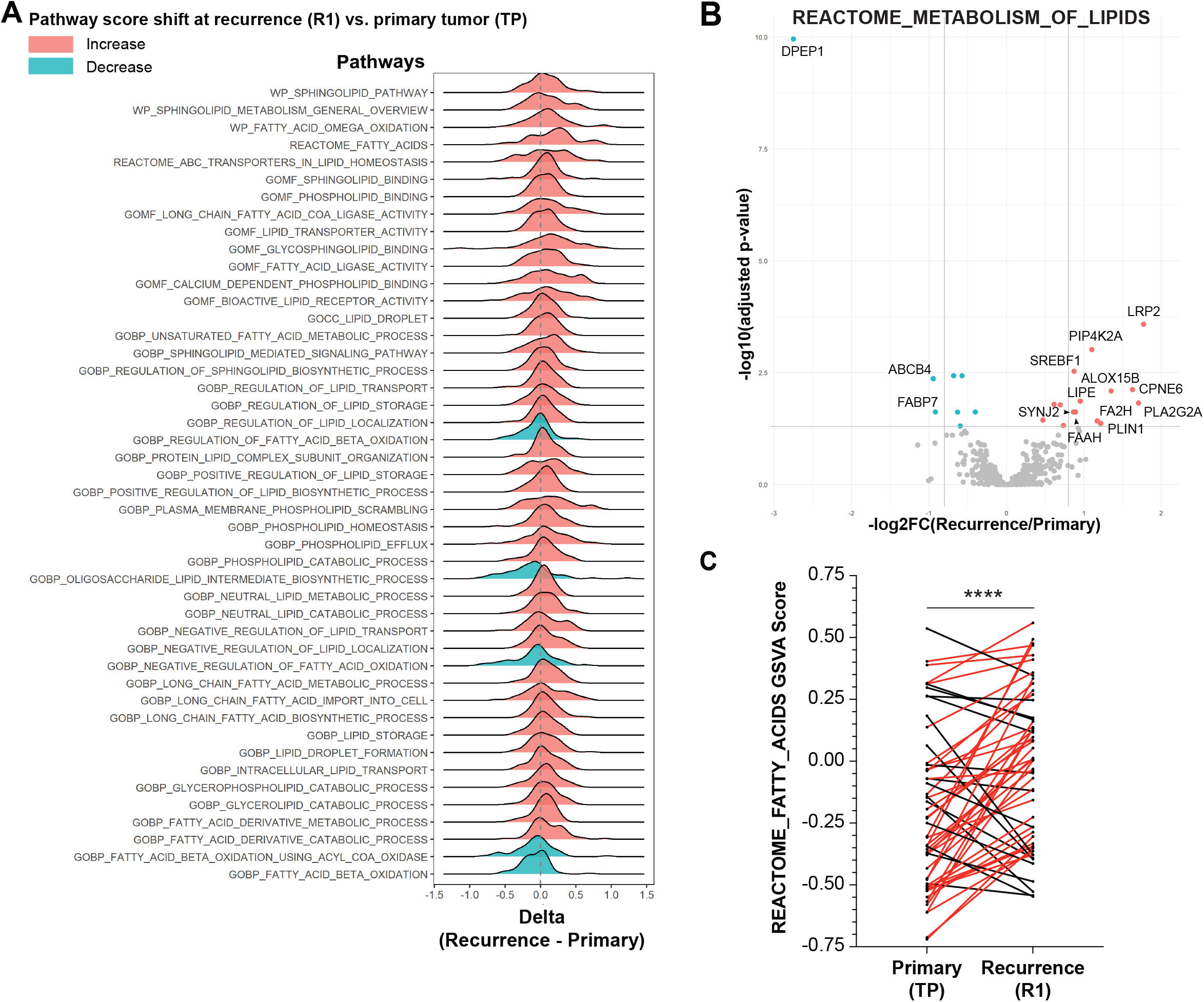
Lipid metabolism is enriched in recurrent glioblastoma tumors. (**A**) Ridge plot shows the distribution of delta values for each of the most significant (p<0.05) pathways by paired t-test of GSVA scores between paired primary and recurrent GBM tumors. Red indicates pathways whose scores increased overall between primary and recurrence (average delta value > 0, n = 41). Blue indicates pathways whose scores decreased overall between primary and recurrence (average delta < 0, n = 5). (**B**) Volcano plot shows differential expression using REACTOME_METABOLISM_OF_LIPIDS in paired recurrent versus primary human Glioblastoma tumors. The two vertical grey lines delineate a log2 fold change of > 0.8 or < -0.8. The horizontal grey line represents an adjusted p-value of < 0.05 after Benjamini-Hochberg correction. The labelled genes meet each of these cutoffs (adjusted p-value < 0.05, and log2 fold change > |0.8|). (**C**) REACTOME_FATTY_ACIDS GSVA scores between paired primary and recurrent GBM tumors. Red lines represent an increase in score and black lines represent a decrease in score in recurrent versus primary for one patient. ****p<0.0001, using Wilcoxon signed-rank test.

## RESULTS

### RT enhances bioenergetic capacity and fuels flexibility of irradiated GBM cells

Mitochondria can adapt in response to cues from the TME to sustain cellular energy in the form of ATP produced by oxidative phosphorylation (OXPHOS). The bioenergetic parameter of mitochondrial fitness is the spare respiratory capacity (SRC), which is the extra mitochondrial capacity available in a cell to produce energy under genotoxic stressors such as RT. To explore whether RT impacts the bioenergetic demand of GBM, we first used Seahorse XF Mito Stress assays to evaluate mitochondrial function in two murine GBM cell lines (GL261 and CT2A). Irradiation of GBM cells significantly increased basal and maximal oxygen consumption rate (OCR) and extra cellular acidification rates (ECAR) indicative of enhanced mitochondrial respiration and glycolysis of GBM cells post 10 Gy RT (Fig. 1A and 1B). As anticipated, these observations resulted in a significant increase of ATP and SCR in irradiated GBM cells.

To identify which oxidative pathway was responsible for the enhanced basal energetic demands in irradiated GBM cells, we investigated the relative contributions of glucose, glutamine and long-chain fatty acid oxidation (FAO) to basal respiration by Seahorse XF Mito Fuel Flex assays. As expected, both GL261 and CT2A mainly rely on glucose consumption as a mitochondrial fuel (Fig. 1C and 1D). Surprinsingly, RT did significantly enhance the contribution of glucose in GL261 but not in CT2A. Interestingly, the ability to consume glutamine and fatty acids in basal respiration was increased in both cell line 24hrs after RT (Fig. 1C and 1D). Of note, in both cell lines, FAO was the most altered with up to a 3-fold increase post RT (Fig. 1C and 1D). These data demonstrate that irradiation of GBM cells is associated with a greater bioenergetic capacity and fuel flexibility.

### RT increases mitochondrial activity to use alternative pathways as energy sources

To further investigate RT-induced metabolic changes in GBM cells, targeted detection of metabolites belonging to glycolysis, tricarboxylic acid (TCA) intermediates and amino acids were performed in GL261 cells 24hrs post 0, 6 and 10 Gy irradiation. Unsupervised principal component analysis (PCA) revealed that replicates from the same group were closely related and can be distinguished from other radiation groups (Fig. 1E), with principal component 1 (PC1) resolving 80.7% of the variability. Based on the degree of similarity of metabolite abundance profiles, hierarchical clustering analysis was performed to show the global view of all metabolites detected in GL261 across radiation groups using MetaboAnalyst 5.0 software. The heatmap revealed that 64 metabolites were differentially represented with a dose-dependent increase in the intensity of most metabolites (Fig. 1F and Supplementary Table 1). Metabolite set enrichment analysis was then performed to identify the most enriched metabolic pathways in GL261 cells. As expected, pathways that reflect DNA damage (27–29), namely the aminoacyl-tRNA biosynthesis, pentose phosphate and nicotinamide, were showed high levels of enrichment (Fig. 1G). Importantly, consistent with our previous data (Fig. 1C), the glutamine metabolism was significantly enriched in response to RT (Fig. 1G).

Altogether, these results show that RT increases mitochondrial activity and the ability to use alternative pathways as energy sources such as glutamine and FAO.

### RT perturbs the metabolism of GBM to promote the generation of unsaturated fatty acids

Because glutamine is a well-known carbon source for *de novo* lipogenesis (30) and RT increases the contribution of fatty acids (FAs) to cope with energy demand (Fig. 1C and 1D), we determined the free fatty acid profiles 24hrs post RT in irradiated GBM cells (Fig. 2 and Supplementary Table 2). Out of the 40 fatty acids we probed, all of them were represented in GBM cells with a baseline level more pronounced in GL261 as compared to CT2A (Supplementary Fig. 1). Interestingly, RT significantly increased the overall content of all FAs. However, unsaturated fatty acids (UFAs) were the most altered in response to genotoxic stress from RT. Specifically, our data shows that RT significantly promotes UFAs such as oleic acid (C18:1), linoleic acid (C18:2), arachidonic acid (AA; C20:4), eicosapentaenoic acid (EPA; C20:5) and docohexaenoic acid (DHA; C22:6) (Fig. 2B and 2D).

The magnitude of metabolic changes may be linked to variables pertaining to the TME including, but not limited to, hypoxia and the availability of nutrients. In that context, to confirm that RT was altering GBM metabolism towards the generation of UFAs *in vivo*, we performed mass spectrometry imaging of brain sections bearing CT2A tumors. To that end, C57BL/6N mice bearing intracranial GBM received tumor targeted irradiation given in a single fraction of 10 Gy on day 10. Brains were harvested on day 15 for spatial metabolomic analysis. Adjacent brain tumor sections were stained with H&E to delinate the tumor area (or region of interet; ROI). The mean abundance of each metabolite of the ROI was then determined (Fig. 3A). Unsupervised PCA showed that replicates from the same group were closely clustered and that irradiated CT2A tumors can be distinguished from control (PC1 90.2%, Fig. 3B). The heat map indicated that 103 metabolites were significantly differentially altered in irradiated CT2A tumors (Fig. 3C and Supplementary Table 3) and confirmed the increase of metabolites linked to glycolysis, pentose phosphate and TCA cycle (e.g. aconitic acid and succinic acid). Additionally, supporting our *in vitro* data, results indicate that RT enhanced glutamine and lipid metabolism with increased levels of metabolites belonging to glutamine/glutamate (e.g. glutamine and glutamic acid) and free fatty acid pathways (Fig. 3C and 3D). Specifically, further validating our *in vitro* data, several UFAs were significantly augmented in irradiated CT2A tumors, including DHA, AA and lysophosphatidic acid (LPA; C22:6) (Fig. 3D). Of notice, even though some glycerolipids were decreased in response to RT, the majority were over-represented in CT2A tumors (Fig. 3C and 3D).

Importantly, no significant augmentation of metabolites was observed in non-tumor areas from matched anatomical regions, thus confirming the selective increase of UFAs in irradiated CT2A tumors (Supplementary Fig. 2 and Supplementary Table 4).

Overall, these results show that tumor irradiation promotes a cellular pool of UFAs that can later serve as building blocks for glycerolipids such as phosphatidic acid (PA), phosphatidylserines (PS), phosphoethanolamines (PE), sphingomyelin and eicosanoids.

### Accumulation of FAs promotes lipid droplets in irradiated GBM cells

Excess of FAs induces lipotoxicity that is associated with endoplasmic reticulum (ER) stress, mitochondrial dysfunction and cell death (31). To prevent this from occurring, cancer cells may generate lipid droplets (LD), a lipid-rich organelle. LDs store neutral lipids, mostly triacylglycerol (TAG) and cholesteryl esters, and balance FA uptake, storage and use according to the energy demands. It has been demonstrated that LDs act as transient buffers that reduce FA lipotoxicity (32). In that context, we asked whether RT-induced FAs result in the formation of LDs to counteract lipotoxicity and support GBM survival in response to RT. To test this hypothesis, we subjected two murine (GL261 and CT2A) and one human (U118) GBM cell lines to a radiation dose escalation and determined the accumulation of neutral lipids by BODIPY and LipidTOX stainings. We found that irradiation of GBM cells results in a significant increase of lipids in a dose-dependent fashion (Fig. 4A and Supplementary Fig. 3). Visualization of LDs in irradiated murine GBM cells confirmed the enhanced formation of LDs 24hrs post RT (Fig. 4B). To evaluate whether this effect was maintained over time, we assessed the accumulation of neutral lipids up to 7 days after RT. Our data demonstrated a sustained increase of lipids in irradiated GBM cells, an effect that is magnified over time in all the cell lines we tested (Fig. 4C).

Because LDs are major sites for prostaglandin E2 (PGE2) synthesis (33), we asked whether RT was promoting the secretion of PGE2 in GBM cells. Consistent with the formation of LDs in irradiated GBM cells, we found a significant augmentation and a trend of increasing PGE2 production 24 hrs post RT in CT2A (*p=0.0136) and GL261 (p=0.0693) cells respectively (Fig. 4D).

Overall, these findings show that RT induces neutral lipid accumulation and storage, an effect that persists for at least 7 days *in vitro* and leads to the secretion of PGE2 by irradiated GBM cells.

### Radiation-induced FASN is responsible for the accumulation of lipids in irradiated GBM

Because FASN is responsible for the *de novo* production of most (if not all) unsaturated lipids (Fig. 4E) (34), we asked whether FASN modulates the synthesis of lipids post-RT. We first irradiated GL261 and CT2A and determined the protein expression of FASN by immunofluorescence. Our data showed an upregulation of FASN in GBM cells 24hrs post RT (Fig. 4F). To interrogate whether FASN was responsible for the accumulation of lipids in irradiated GBM cells, we generated CRISPR/Cas9 engineered sgControl (control) or sgFasn (Fasn inactivation) CT2A and GL261 cells. FASN deficiency was validated by western blot (Supplementary Fig. 4A).

In line with our previous data, irradiated but not control GBM cells accumulated neutral lipids in both GL261 and CT2A. Interestingly, confirming the role of FASN in the RT-induced generation of lipids, lipid accumulation was significantly reduced in FASN-deficient GBM cells (Fig. 4G) or in GBM cells treated with FASN inhibitors (Fig. 4H). Of note, the decrease of neutral lipid accumulation was not complete in GBM-sgFasn cells or post FASN inhibition, suggesting that another pathway might promote lipid accumulation in irradiated GBM cells (Fig. 4G-4H).

### Lipid accumulation copes for ER stress and limits apoptosis of irradiated GBM

Reports indicate that excessive synthesis of lipids is an adaptive mechanism of ER to cope with cell stress and avoid apoptosis (35–37). Our results indicate that RT is eliciting a lipogenic phenotype in GBM (Figs. 2-4). Thus, we asked whether FASN-mediated lipid synthesis is a resistance mechanism that maintains survival and resolves ER stress in irradiated GBM cells. To test this hypothesis, we used CRISPR/Cas9 and shRNA-mediated FASN knockdown (KD) cells. GL261 and CT2A derivatives with selective KD of FASN using a doxycycline (DOX) inducible construct (Supplementary Fig. 4B). The shRNA was introduced into a pTRIPZ inducible lentiviral shRNA vector containing the tetracycline response element (TRE) promoter, which leads to shRNA expression in the presence of DOX.

Having confirmed the KD of FASN in GBM cells (Supplementary Fig. 4C), we investigated the role of FASN-mediated lipid synthesis and ER stress in irradiated GBM cells. Hallmarks of ER stress are, but not limited to, loss of mitochondrial integrity and the activation of the unfolded protein response (UPR) pathway (38). We first analyzed mitochondria by transmission electronic microscopy (TEM) in GL261 and CT2A derivatives 24hrs post RT. Our data revealed a loss of mitochondrial shape and a decrease of their crypts in FASN-deficient GBM cells, indicative of a mitochondrial dysfunction. Interestingly, irradiation did not enhance this effect, thus suggesting that the absence of FASN alone can perturb the energy supply of GBM cells at least in FASN KD cells (Fig. 5A). Further supporting the role of RT-induced FASN lipid synthesis in preventing mitochondrial stress, inhibition of FASN significantly reduced the increase of maximal respiration and SRC of irradiated GBM cells (Fig. 5B).

The UPR pathway is triggered by the onset of ER stress to restore cell homeostasis (39). There are three primary branches of the UPR pathway and each is mediated by distinct ER proteins namely inositol-requiring enzyme 1 (IRE1), protein kinase R-like endoplasmic reticulum kinase (PERK) and activating transcription factor 6 (ATF6) (39). During ER stress, the ER chaperone BiP (immunoglobin binding protein) dissociates from these ER stress transducers leading to their activation. If ER protein homeostasis is not resolved, the prolonged activation of UPR may initiate apoptosis via the upregulation of the C/EBP Homologous Protein (CHOP). Therefore, to interrogate whether RT-induced FASN is controlling ER stress, we determined the expression of BiP and CHOP 24 hrs post RT in GBM-sgFasn cells. Tunicamycin (2 μg/mL) was added to GBM cells to induce ER stress as a positive control. A slight baseline protein expression of BiP and CHOP was observed in untreated or irradiated GBM cells. However, the absence of FASN markedly increased BiP and CHOP in untreated GBM cells, an effect that was amplified in irradiated GBM (Supplementary Fig. 5A).

Although evidence indicates that FASN can resolve ER stress of irradiated GBM cells, it is still unclear whether it protects them from cell death. To address this question, we first measured the extracellular release of High mobility group box protein 1 (HMGB1), a damage-associated molecular pattern (DAMP) that is emitted by dying cells, 24 hours post RT. As expected, RT increases the release of HMGB1 in both GBM cells. FASN KD promoted extracellular accumulation of HMGB1 in GL261 but not in CT2A cells. Interestingly, HMGB1 release was enhanced in irradiated FASN-deficient GL261 and CT2A cells (Supplementary Fig. 5B), suggesting that cytocidal properties of RT might be improved when FASN is impaired.

To test this hypothesis, we first assessed the survival ability of irradiated cells in the presence of a FASN inhibitor (FASNi). As expected, 4 Gy irradiation impacted the clonogenic survival of GL261 and CT2A cells. While the presence of FASNi did not affect the formation of GBM clones, inhibition of FASN significantly enhanced radiosensitivity in both GBM cell lines (Fig. 5C), thus suggesting that radiation-induced FASN lipid synthesis is a mechanism of radioresistance that prevent cells from undergoing apoptosis. To confirm this concept, we determined apoptosis and necrosis 24hrs after irradiation by flowcytometry. We found that apoptosis and necrosis were greater in GBM-sgFasn cells as compared to sg-Control cells. Interestingly, RT significantly increased apoptosis but did not further promote necrosis in both FASN deficient GBM cells (Fig. 5D). Supporting these observations, similar results were obtained with a FASNi (Fig. 5E).

Overall, these results indicate that FASN-mediated lipid accumulation prevents ER stress and evades apoptosis to cope with genotoxic insult and promote survival of irradiated GBM cells.

### Combination of FASN blockade with focal radiation therapy prolongs survival of tumor bearing mice

To determine whether combination of irradiation with FASN blockade is associated with therapeutic activity, C57BL/N mice bearing syngeneic CT2A intracranial tumors received locoregional infusion of a FASN inhibitor from day 10 to day 24. Mice received tumor-targeted irradiation in a single fraction of 10 Gy starting day 10 and were followed for survival (Fig. 5F). FASN blockade as monotherapy did not have any effect on survival (ns; p=0.0926). As expected, RT significantly increased survival of GBM-tumor bearing animals (**p=0.0091 versus control). However, combination of RT with FASN blockade significantly doubled the median overall survival of GBM-bearing animals when compared to control mice (**p=0.0015 versus control; Fig. 5F); demonstrating the therapeutic relevance of our *in vitro* findings.

### Lipid metabolism gene signature is enriched in recurrent GBM patients

To test the translational value of our findings, we performed analysis on paired RNAseq files from human GBM patients included in the GLASS Consortium (23). We hypothesized that GBM patients at their first recurrence would exhibit increased lipid and fatty acid metabolism signatures as a response to RT treatment. To address this question, we compared matched primary tumor (TP) to first recurrence (R1) from 55 GBM patients that received at least 50 Gy of RT between the two time-points (Supplementary Table 5). We used GSVA to score each of these samples using 193 lipid and fatty-acid related pathways from the Molecular Signature Database (MSigDB) (25). We found 46 pathways with significant (paired t-test p<0.05) score changes between TP and R1; 41 which were upregulated at recurrence and 5 which were downregulated at recurrence (Fig. 6A; Supplementary Table 6).

We then performed differential gene expression analysis between R1 and TP tumors using expression counts of 712 genes included in the Metabolism of Lipids pathway from REACTOME (Supplementary Table 7). We found increased expression of genes envolved in LD formation (LIPE logFold Change (FC) = 0.954, p = 0.0136; PLIN1 FC = 1.215, p = 0.0427), in FA synthesis (SREBF1 FC = 0.873, p = 0.0029; PLAG2G2A FC = 1.705, p = 0.0150) and in the arachidonic acid metabolism (FAAH FC = 0.892, p = 0.0238; ALOX15B FC = 1.351, p = 0.0081) (Fig. 6B, Supplementary Table 5). This is consistent with our data that indicate an increase of fatty acid synthesis and UFAs (namely AA, EPA and DHA) (Figs. 2-3) and an accumulation of neutral lipids and LD post RT (Fig. 4).

To further evaluate fatty acid metabolism per individual patients, we calculate the GSVA enrichment score using expression counts of the genes included in the Fatty Acid pathway from REACTOME. Supporting our findings, we found a significant upregulation of the fatty acid pathway score in 39 patients out of 55 at recurrence (****p<0.0001; Fig. 6C and Supplementary Table 8).

Overall, these data are consistent with our preclinical findings and supports the relevance of targeting lipid metabolism in human GBM to decrease resistance to RT and limit GBM recurrence.

## DISCUSSION

RT is the standard-of-care for the management of GBM and is the most widely used treatment for inoperable brain tumors. However, GBM relapses often arise within the irradiated margins (7), suggesting that the TME of GBM post RT generates resistance mechanisms that may support GBM recurrence. The mechanisms by which RT drives GBM regrowth remain elusive.

Here, we showed that RT provides metabolic plasticity for GBM to survive genotoxic insults and acquire adaptive resistance mechanisms. Specifically, we identified the biosynthesis of fatty acids as a dominant metabolic pathway that protects irradiated GBM cells from undergoing ER stress; thereby permitting GBM survival and the escape of RT-induced apoptosis. These results have important implications for the use of lipid metabolism targeting agents in patients, and suggest that combination of RT with such compounds might be required to develop treatments that are more effective and resistant to GBM recurrence.

Prior research conducted in pancreatic tumors demonstrated that mitochondrial activity and OXPHOS are activated in resistant tumors to survive anti-cancer modalities such as RT (40). Supporting this notion, our findings demonstrate that mitochondria activate OXPHOS to generate lipids and cope with the energy demand post RT. Specifically, the ability to use FAO for basal respiration was increased in both of our murine GBM cell models despite the fact that they displayed disctinct metabolic profiles at baseline. These results are in line with a previous study in which FAO was identified as a dominant metabolic program that provides GBM metabolic flexibility to adapt to the nutrient heterogeneity of its TME (41).

Surprinsingly, our results show that RT increases both FAO and fatty acid synthesis (FAS) *in vitro*. While FAO and FAS are usually considered incompatible due to the inhibition of malonyl-CoA, accumulating evidence support the hypothesis of both processes coexisting and feeding each other in some cancer cells that overproduced NADPH in response to oxidative stress such as breast cancer (42). Whether this phenomenon is occuring in our preclinical models of GBM remains to be investigated.

Additionally, we found that irradiated GBM (both *in vitro* and *in vivo*) rewires its energy supply towards the synthesis of UFAs to presumably generate a cellular pool of fatty acids that can be utilized as building blocks for complex lipids involved in structure, fluidity and function of the cellular membrane. Similar findings have been reported in the field of radiation countermeasures where a specific lipidomic signature (mostly belonging to PUFAs) was identified in serum of healthy mice exposed to a radiation dose of 8 Gy (43).

Importantly, our findings demonstrate that accumulation of UFAs in response to RT leads to the formation of LDs, which are major sites for PGE2 synthesis and have been ascribed roles in tumor cell proliferation, cancer stemness and immunosuppression (44–46). Our *in silico* analysis of recurrent GBM patients confirms the increase of arachidonic metabolism with significant upregulation of arachidonate 15 lipoxygenase type B (ALOX15B), thus underscoring that RT might promote the formation of UFAs to synthesize eicosanoids, and more specifically the prostanoid family of lipids in recurrent GBM patients.

Our study suggests that the biological implications of lipid metabolism reprogramming subsequent to tumor targeted irradiation are to protect GBM from apoptosis by resolving ER stress. Consequently, RT-induced lipid metabolism may drive resistance of GBM. With limitations pertaining to retrospective *in silico* studies, our findings support this concept with a significant enrichment of fatty acid in recurrent GBM patients. Additionally, while the clinical relevance of our findings need to be validated, our study suggests that combination of lipid metabolism inhibitors with ER stress inducers may represent a novel strategy to prolong survival of irradiated GBM patients.

PGE2 is known to play a role in regulating inflammation that drives cancer initiation and progression. Several studies have shown that PGE2 can activate growth factor signaling, resistance to apoptosis and enhance immune evasion (47). PGE2 is the major product of the cyclooxygenases (COXs). While COX-1 is constitutively at basal levels of many cells, COX-2 is induced in response to several stimuli to mediate pathological events that are associated with inflammatory processes (48). In fact, GBM highly expresses COX-2 which led to the development of COX-2 inhibitors such as celecoxib that is well-known to induce ER stress. However, in a randomized phase II clinical trial with newly diagnosed GBM patients, RT+temozolomide (TMZ) combined with celecoxib failed to improve patients’ overall survival (NCT00112502, (49)). Based on our data, it is plausible that celecoxib was unable to induce ER stress due to RT-induced FASN metabolism in these patients. Further, dual blockade of celecoxib with FASN inhibition promotes ER stress and subsequent cell death in mucinous colon cancer (50). As FASN inhibitors are actively tested in clinical trials for multiple tumors including GBM (NCT03808558, NCT02980029, NCT03179904, NCT03032484), this work supports the use of these compounds to enhance the efficacy of ER stress inducers as well as COX-2 inhibitors to potentially prolong the survival of GBM patients.

On another note, PGE2, has also been reported to promote an immunosuppressive phenotype of tumor-associated macrophages (TAMs) by initiating a signal transduction cascade to increase the programmed death-ligand 1 (PD-L1) transcriptional levels of TAMs (46). Our work shows that (i) arachidonic acid, a precursor for PEG2 synthesis, is more abundant post RT, (ii) PGE2 is upregulated in irradiated GBM cells and (iii) arachidonic metabolism is upregulated in recurrent GBM patients. Thus, it is conceivable that FASN upregulation in response to RT is a major obstacle to the efficacy of anti-programmed death-1 (PD-1) in GBM. Previous clinical trials targeting PD-1 alone led to only 8% of objective response rate in recurrent GBM (Checkmate 143) (51). As all patients from Checkmate 143 received prior RT and TMZ, it is possible that RT-induced FASN may have driven PGE2-mediated immunosuppression to limit anti-PD-1 efficacy in these patients. While further evidence are warranted, our work also support the combination of FASN inhibitors with PD-1 blockade to improve the outcome of GBM patients in the context of RT.

## CONCLUSIONS

In conclusion, our results identify the role of lipids in providing metabolic plasticity of irradiated GBM, therefore allowing GBM cells to accommodate to genotoxic stress and develop resistance mechanisms by at least icreasing UFAs to synthesize eicosanoids, and more specifically the prostanoid family. Overall these data underscore the role of UFAs in shaping the TME and cancer progression of irradiated GBM. Although the role of UFAs in RT-induced resistance of GBM remains to be confirmed, our data provide the rationale for testing UFA targeting agents in the clinic, and highlight a potential new actionable target to improve response in GBM patients post RT.

## Acknowledgments

We are acknowledging the Weill Cornell Medicine Proteomics and Metabolomics Core Facility for their assistance in profiling free fatty acids in irradiated murine glioblastoma cells. We are grateful to the Electron Microscopy & Histology services of the Weill Cornell Medicine Microscopy & Image Analysis Core for their assistance in transmission electronic images. We thank the Weill Cornell Advanced Biomolecular Analysis Core (ABAC) for MALDI-MS sample and data processing. We are acknowledging Dr. Manuela Buonanno from Columbia University, Department of Radiation Oncology for her advice pertaining to the clonogenic assay.

## FIGURE CAPTIONS

**Supplementary Figure 1: Comparison of fatty acids baseline content in glioblastoma cells.** Differentially modulated fatty acids determined by free fatty acid profiling by mass spectrometry in GL261 vs CT2A cells. Experiment was done in duplicate with n=5 per group.

**Supplementary Figure 2: Quantification of metabolites analyzed by MALDI imaging in non-tumor areas.** H&E of brain sections from animals in Figure 3. Scale 5mm. Metabolites were quantified in the non-tumor area delineated with black lines in the H&E sections. Peak intensity and representative images of metabolites. Values represent mean ± SEM. Each dot represents one animal. ns, not significant; *p<0.05, using unpaired Student’s t test.

**Supplementary Figure 3: Radiation induces accumulation of lipids in glioblastoma cells**. GL261 and CT2A cells were subjected to different radiation doses from 2 Gy to 10 Gy *in vitro* and 24hrs later, neutral lipids accumulation was determined by LipidTOX staining and flow cytometry.

**Supplementary Figure 4: Characterization of genetically modified glioblastoma cells.** (**A**) Western blot analysis of FASN protein expression in GL261 and CT2A CRISPR/Cas9 sgControl or sgFasn clones, obtained with 2 different anti-Fasn guide RNAs. (**B**) Modified pTRIPZ lentiviral vector with tetracycline-inducible shRNA directed against the mouse *Fasn* gene (shFASN) or its non-silencing control (shNS). Fasn knockdown in GL261 and CT2A cells was confirmed by western blot 10 days after doxycycline induction (**C**).

**Supplementary Figure 5: Radiation-induced FASN prevents ER stress and apoptosis in irradiated GBM.** (**A**) Western blot results showing expression of BIP and CHOP, in CRISPR/Cas9 sgControl or sgFasn CT2A cells 24hrs after 10 Gy irradiation or sgControl cells treated for 24hrs with tunicamycin 2μg/ml. VINCULIN expression levels were used as a loading control. (**B**) HMGB1 secretion was determined by ELISA 24hrs after 10 Gy irradiation of shNon-silencing (shNS) or shFASN GL261 and CT2A cells. Results are expressed in pg/mL ± SEM for 100,000 live cells. Values represent mean ± SEM. Each dot represents one replicate. ns, not significant; ****p<0.0001, using one-way ANOVA followed by Tukey post hoc comparisons (B). Experiment was done in duplicate with n=3 per group.

**Supplementary Table 1: Peak intensities of polar metabolites in irradiated GL261.**

**Supplementary Table 2: Peak intensities of free fatty acids in irradiated GL261 and CT2A.**

**Supplementary Table 3: Average intensities (peak area) of metabolites from CT2A tumors.**

**Supplementary Table 4: Average intensities (peak area) of metabolites from corresponding non-tumor regions.**

**Supplementary Table 5: Clinical data of GBM patients from the GLASS consortium dataset**

**Supplementary Table 6: Differential gene expression of all lipids and fatty acids pathways in Recurrent versus Primary GBM patients.**

**Supplementary Table 8: Differential gene expression of metabolism of fatty acids pathway in Recurrent versus Primary GBM patients.**

