## Supplementary figures and images for "Radiation therapy promotes unsaturated fatty acids to maintain survival of glioblastoma"

### Supplementary Figure 1

#### Supplementary Figure 1

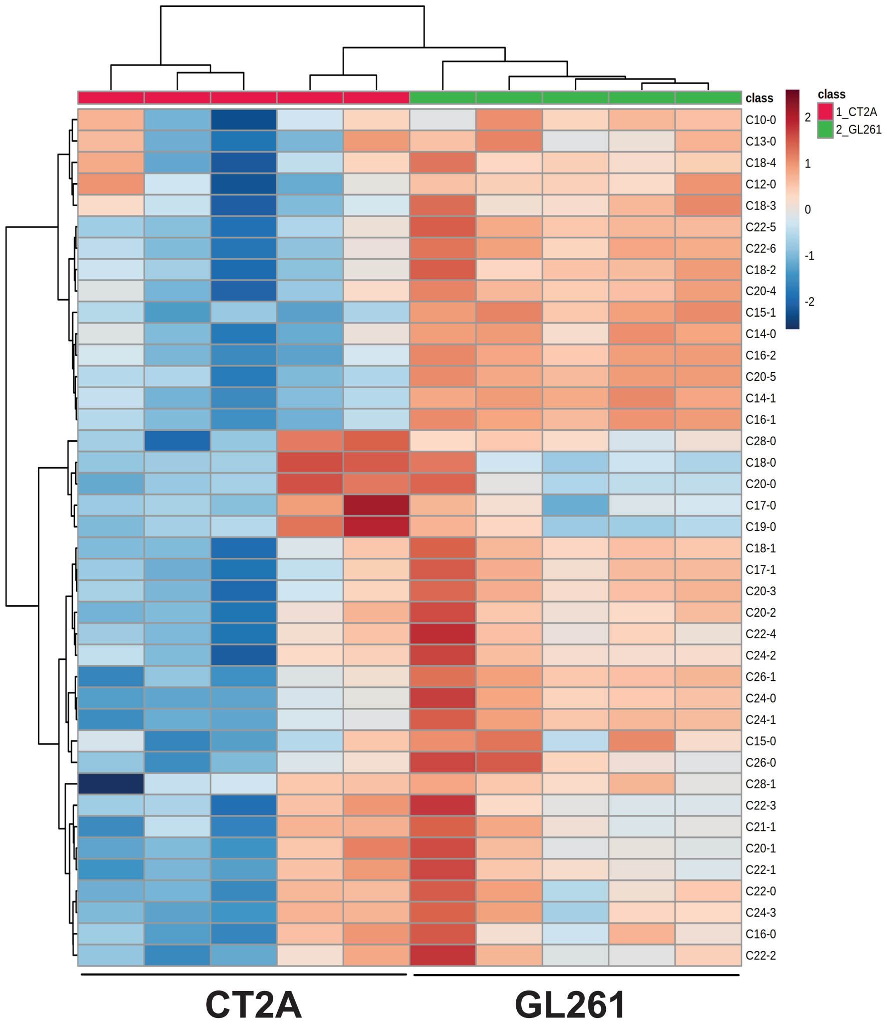

### Supplementary Figure 2

#### Supplementary Figure 2

A

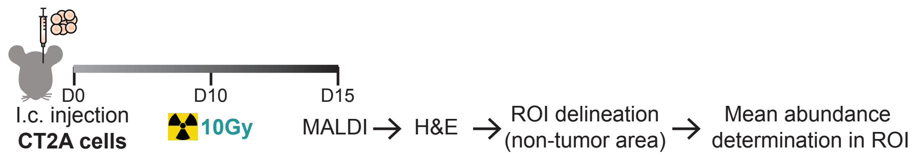


B

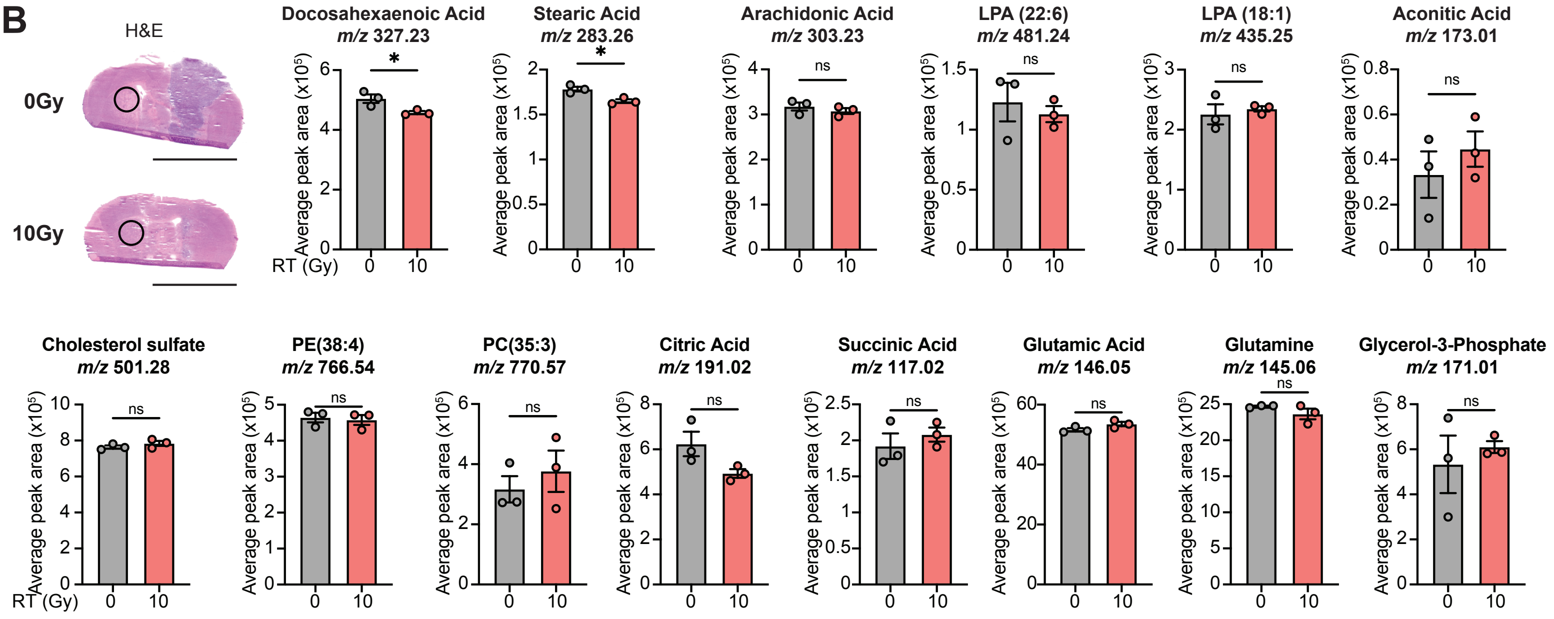

### Supplementary Figure 3

#### Supplementary Figure 3

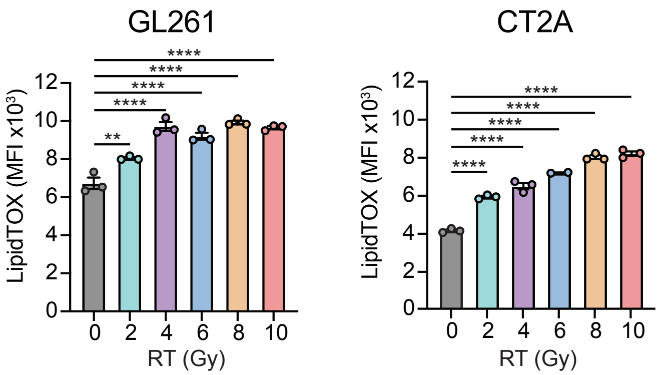

### Supplementary Figure 4

##### Supplementary Figure 4

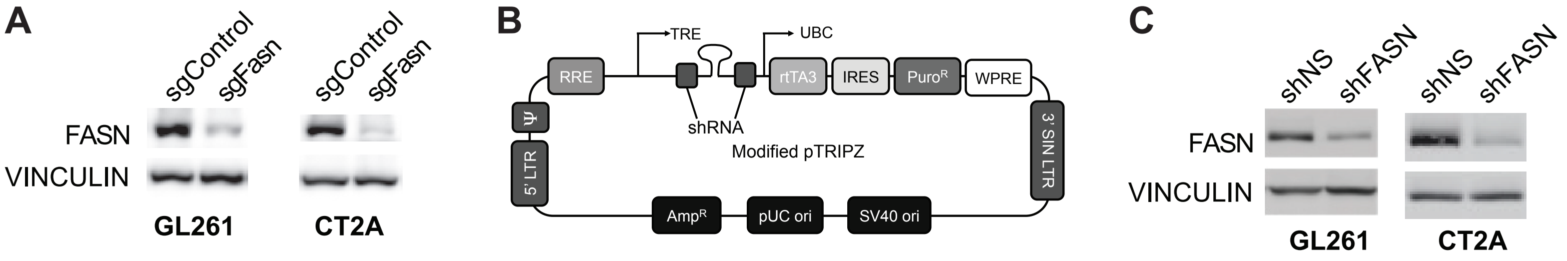

### Supplementary Figure 5

#### Supplementary Figure 5

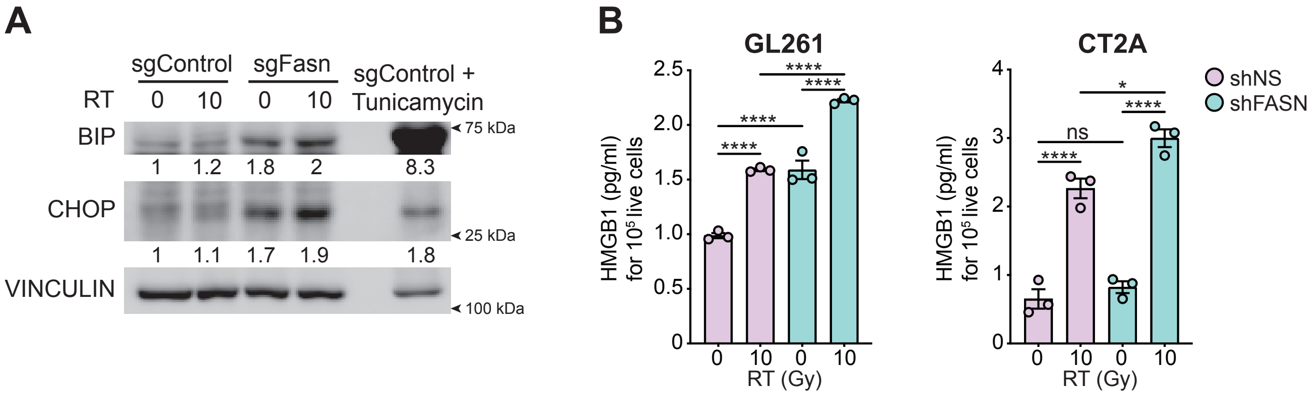
